# Enhanced Ca^2+^-Driven Arrhythmias in Female Patients with Atrial Fibrillation: Insights from Computational Modeling

**DOI:** 10.1101/2024.03.04.583217

**Authors:** Xianwei Zhang, Yixuan Wu, Charlotte Smith, William E. Louch, Stefano Morotti, Dobromir Dobrev, Eleonora Grandi, Haibo Ni

**Author notes:** Correspondence: Eleonora Grandi, Department of Pharmacology, University of California Davis 451 Health Science Drive, GBSF 3503, Davis, CA 95616, USA, Haibo Ni, Department of Pharmacology, University of California Davis 451 Health Science Drive, GBSF 3503, Davis, CA 95616, USA. Contributed Equally to the work (Xianwei Zhang, Yixuan Wu). Shared senior authorship (Eleonora Grandi, Haibo Ni).

## Abstract

**Background and Aims:** Substantial sex-based differences have been reported in atrial fibrillation (AF), with female patients experiencing worse symptoms, increased complications from drug side effects or ablation, and elevated risk of AF-related stroke and mortality. Recent studies revealed sex-specific alterations in AF-associated Ca^2+^ dysregulation, whereby female cardiomyocytes more frequently exhibit potentially proarrhythmic Ca^2+^-driven instabilities compared to male cardiomyocytes. In this study, we aim to gain a mechanistic understanding of the Ca^2+^-handling disturbances and Ca^2+^-driven arrhythmogenic events in males vs females and establish their responses to Ca^2+^-targeted interventions.

**Methods and Results:** We incorporated known sex differences and AF-associated changes in the expression and phosphorylation of key Ca^2+^-handling proteins and in ultrastructural properties and dimensions of atrial cardiomyocytes into our recently developed 3D atrial cardiomyocyte model that couples electrophysiology with spatially detailed Ca^2+^-handling processes. Our simulations of quiescent cardiomyocytes show increased incidence of Ca^2+^ sparks in female vs male myocytes in AF, in agreement with previous experimental reports. Additionally, our female model exhibited elevated propensity to develop pacing-induced spontaneous Ca^2+^ releases (SCRs) and augmented beat-to-beat variability in action potential (AP)-elicited Ca^2+^ transients compared with the male model. Parameter sensitivity analysis uncovered precise arrhythmogenic contributions of each component that was implicated in sex and/or AF alterations. Specifically, increased ryanodine receptor phosphorylation in female AF cardiomyocytes emerged as the major SCR contributor, while reduced L-type Ca^2+^ current was protective against SCRs for male AF cardiomyocytes. Furthermore, simulations of tentative Ca^2+^-targeted interventions identified potential strategies to attenuate Ca^2+^-driven arrhythmogenic events in female atria (e.g., t-tubule restoration, and inhibition of ryanodine receptor and sarcoplasmic/endoplasmic reticulum Ca²⁺-ATPase), and revealed enhanced efficacy when applied in combination.

**Conclusions:** Our sex-specific computational models of human atrial cardiomyocytes uncover increased propensity to Ca^2+^-driven arrhythmogenic events in female compared to male atrial cardiomyocytes in AF, and point to combined Ca^2+^-targeted interventions as promising approaches to treat AF in female patients. Our study establishes that AF treatment may benefit from sex-dependent strategies informed by sex-specific mechanisms.

**Translational perspective:** Accumulating evidence demonstrates substantial sex-related differences in atrial fibrillation (AF), which is the most common arrhythmia, with female patients faring worse with the condition. By integrating known sex-differential components into our computational atrial cardiomyocyte model we found that female atrial cardiomyocytes in AF exhibit greater propensity to develop Ca^2+^-driven arrhythmia than male cardiomyocytes. Model analyses provided novel mechanistic insights and suggested strategies such as t-tubule restoration, correction of Ca^2+^-handling disturbances, and the combination of both, as promising approaches to treat AF in female patients. Our study uncovers and validate sex-specific AF mechanisms and inform the development of targeted anti-AF strategies.

**Graphical abstract:**
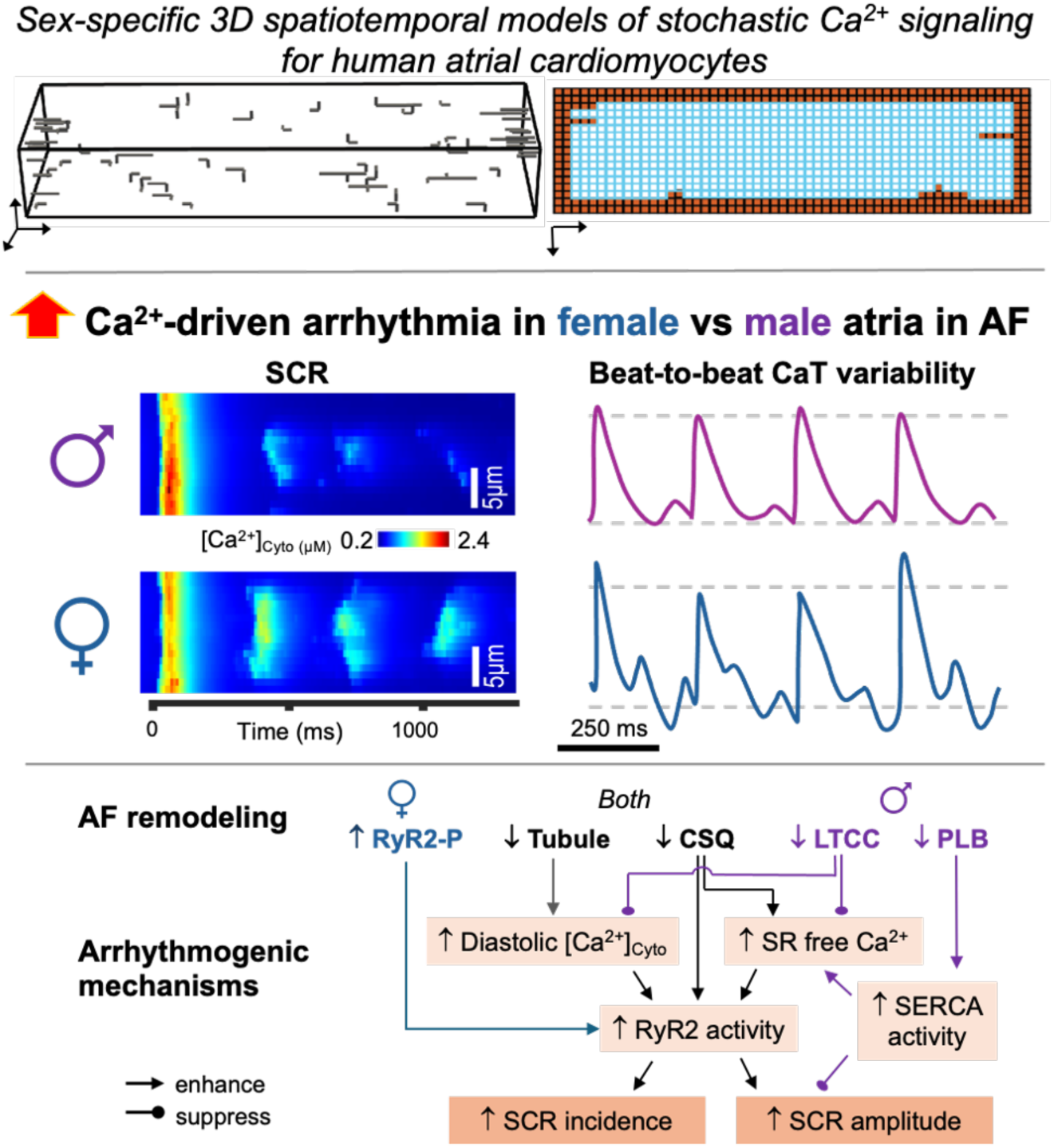
Sex-specific 3D spatiotemporal models of human atrial cardiomyocyte Ca^2+^ signaling reveal a greater propensity to develop Ca^2+^-driven arrhythmic events in female vs male atrial cardiomyocytes in AF. Model analysis links sex-specific AF remodeling to arrhythmogenic mechanisms. AF, atrial fibrillation; SCR, spontaneous Ca^2+^ release; CaT, cytosolic Ca^2+^ transient; RyR2-P, phosphorylated ryanodine receptor type 2 (RyR2); CSQ, calsequestrin; LTCC, L-type Ca^2+^ channel; PLB, phospholamban; SERCA, sarcoendoplasmic reticulum Ca^2+^ ATPase; SR, sarcoplasmic reticulum.

## Introduction

Atrial fibrillation (AF), the world’s most common cardiac arrhythmia, is a global epidemic with a prevalence projected to increase 3-fold in the next 50 years.^1,2^ AF causes significant morbidity and mortality, and places a substantial socioeconomical burden on healthcare systems. Despite decades of advances in both fundamental research and clinical management, current AF therapies (e.g., antiarrhythmic drugs and catheter ablation) exhibit inadequate efficacy^3^ and pose significant risks of adverse effects.^4^ The challenge in developing effective AF therapeutic strategies is exacerbated by the marked heterogeneity in disease phenotypes across patient groups^3^ coupled with a limited understanding of the underlying pathophysiological mechanisms.

Sex-related differences have been reported in the prevalence, clinical manifestation, management strategies, and treatment outcomes of AF.^5–10^ Men develop AF years earlier than women;^5,9,10^ however, women experience worse symptoms, more complications arising from drug side effects or ablation, diminished quality of life, and greater risk of AF-related stroke and mortality.^5,9,10^ Although these differences may be attributed to inherent differences in cardiac function between male and female patients, the insufficient inclusion of female sex in both fundamental research and clinical cohorts^11^ has led to an incomplete understanding of the mechanisms that contribute to these sex differences. Because of this, females often obtain less effective but potentially harmful treatment approaches that lead to worse prognoses. While there is a growing interest in exploring sex differences in ventricular electrophysiology,^12–17^ far less attention has been directed to the mechanistic understanding of the differences between the atria of male and female patients. As such, a systematic investigation of these sex differences is urgently required to discover the precise mechanisms of sex-dependent disparities observed in AF to lay the foundation for defining sex-specific therapeutic strategies.

While AF comprises electrophysiological and structural remodeling, dysregulated Ca^2+^ handling has emerged as a key factor in its pathophysiology.^18–23^ As in the ventricles, Ca^2+^ crucially mediates cardiac excitation-contraction coupling in the atria, whereby electrical excitation of the cardiomyocytes leads to changes in the intracellular Ca^2+^ that mediate mechanical contraction and relaxation by acting on the myofilament machinery.^24^ As a consequence of the coupling between Ca^2+^ and voltage, intracellular Ca^2+^-handling abnormalities in atrial cardiomyocytes from patients with AF^18,19^ not only directly impair contractility,^25^ but could also promote ectopic (triggered) activity that significantly contributes to AF initiation and maintenance.^18,19^ This is driven by spontaneous Ca^2+^ releases (SCRs) from the sarcoplasmic reticulum (SR) during diastole, which cause increased Ca^2+^ extrusion by the electrogenic Na^+^/Ca^2+^ exchanger (NCX) that triggers transmembrane potential afterdepolarizations, leading to ectopic activity in the atria.^18,19,23,26,27^ Additionally, Ca^2+^ dysregulation further promotes electrical and structural remodeling in the atria, thereby increasing the likelihood of AF maintenance.^19^ Therefore, the correction of Ca^2+^-handling abnormalities has emerged as a promising anti-AF strategy.^19,23,28,29^

Previous experimental work has reported sex differences in the expression and phosphorylation of Ca^2+^-handling proteins,^30^ t-tubular system density,^31^ and cell dimensions^31–33^ of atrial cardiomyocytes. For instance, the well-documented AF-typical reduction in L-type Ca^2+^ current (I_CaL_)^34,35^ was observed in atrial cardiomyocytes from male, but not female patients.^30^ Conversely, atrial cardiomyocytes from females with AF displayed a greater incidence of Ca^2+^ sparks than males.^30^ While these findings are indicative of AF-related sex differences in ionic, structural, and functional aspects, their precise arrhythmogenic consequences and sex-specific implications are not yet mechanistically dissected and quantitatively determined. Importantly, there is a scarcity of studies investigating differences in atrial rate dependency of Ca^2+^-handling processes between male and female cardiomyocytes, which are crucial in understanding sex-specific mechanisms of arrhythmogenesis in AF. While experimentally delineating the precise contribution of these aspects to sex differences in AF presents significant challenges, computational models have emerged as a powerful tool for unraveling the subcellular mechanisms of Ca^2+^ dysregulation.^36–39^

In this study, we employed computational modeling to systematically assess the mechanisms of Ca^2+^-dependent arrhythmia in male and female atrial cardiomyocytes. We also compared male vs female responses to various putative therapeutic interventions targeting the subcellular determinants of intracellular Ca^2+^ handling. To achieve this, we utilized our recently developed three-dimensional (3D) human atrial cardiomyocyte model that couples electrophysiology with spatially-detailed subcellular Ca^2+^ signalling.^36,37^ We built a suite of male- and female-specific spatially-detailed models of intracellular Ca^2+^ dynamics in normal sinus rhythm (nSR) and AF conditions by incorporating all known sex- and AF-dependent alterations in Ca^2+^-handling,^30^ t-tubular system density,^31,40^ and cell dimensions.^31–33,41^ We found that female AF atrial cardiomyocytes displayed a higher incidence and amplitude of SCRs, and a greater beat-to-beat instability of the paced Ca^2+^ transients *vs* male, owing mainly to female-specific AF remodeling in ryanodine receptor type-2 (RyR2) phosphorylation. Furthermore, our simulations identified potential intervention strategies targeting the Ca^2+^ handling system that may yield favorable outcomes in female atrial cardiomyocytes. Overall, our study underscores and validates the role of sex-specific differences in AF that warrants future fundamental and clinical studies to develop sex-specific anti-AF approaches.

## Methods

### Baseline model and overall study design

Our recently developed three-dimensional model of the human atrial cardiomyocytes that couples electrophysiology, subcellular Ca^2+^ dynamics, and t-tubular structures^36,37^ was extended to incorporate descriptions of reported subcellular gradients of RyR2 and calsequestrin (CSQ) distribution,^30^ and deemed to represent the male atrial cardiomyocyte in normal sinus rhythm (nSR). This model served as the reference for adjusting parameters to generate additional subcellular Ca^2+^ handling models of the male cardiomyocyte in AF conditions, and of female cardiomyocytes in nSR and AF conditions (**Table 1**, detailed descriptions of parameters are listed in **Supplementary Table S1**). Our sex-specific models were employed to assess the propensity of atrial cardiomyocytes to developing Ca^2+^ sparks in quiescent cells and SCR events following burst-pacing, and to mechanistically investigate the impact of AF- and sex-dependent factors on Ca^2+^-driven arrhythmogenesis. Furthermore, we systematically evaluated the responses of atrial cardiomyocytes to several candidate Ca^2+^-targeted intervention strategies.

**Table 1.**
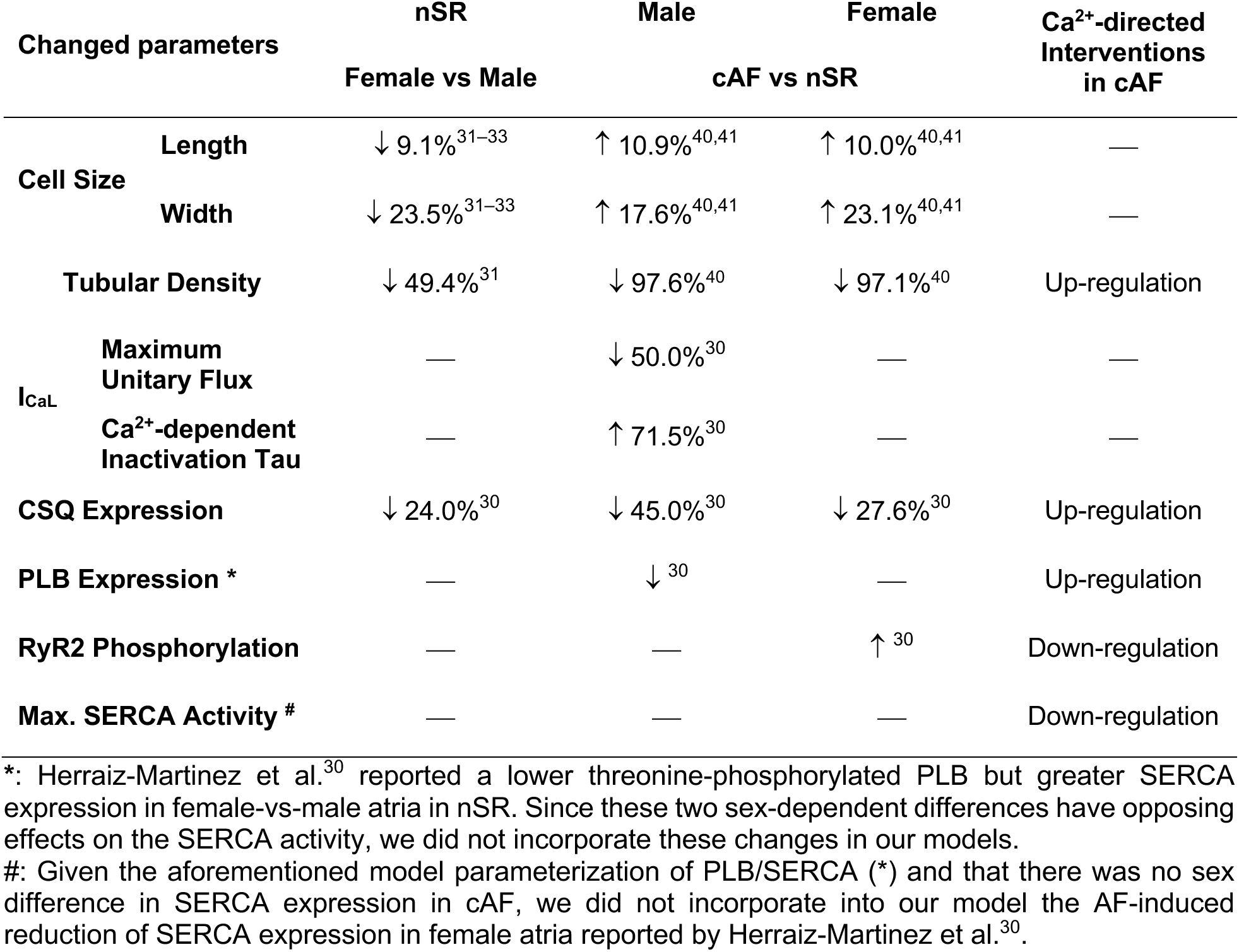
Sex- and AF-dependent changes in the substructure and Ca^2+^-handling processes.

### Updated structural characteristics and subcellular distribution of RyR2s and CSQ in sex-specific atrial cardiomyocyte models

To account for the previously reported heterogeneous subcellular localization of RyR2s and CSQ,^30^ we incorporated the density gradient of RyR2s and CSQ in surface Ca^2+^ release units (CRUs, i.e., the 2 outer CRU layers near cell surface) vs inner (i.e., non-surface) CRUs while keeping the whole-cell total expression unchanged. Specifically, the density ratio of surface-to-inner CRUs was assigned to 1.5 for RyR2s and 3 for CSQ, based on reported subcellular gradients.^30^ We incorporated descriptions of cell dimensions based on experimental findings of cell size differences between sexes and with AF-remodeling, with larger cell dimensions in male vs female cardiomyocytes^31–33^ and in cAF vs nSR conditions^40,41^ (**Table 1**). Additionally, to account for detected differences in t-tubular density between sexes^31^ and AF-remodeling,^40^ the t-tubular density was lower in female vs male cardiomyocyte models, or for cAF vs nSR models (**Table 1**), while maintaining the average branch length of the t-tubular structures. The electrical capacitance of the new sex-specific models of nSR and cAF cardiomyocytes was computed according to the modifications of cell dimension and t-tubular density, as done previously.^36,37^

### Parameterization of Ca^2+^ handling to construct sex-specific atrial cardiomyocyte models

We updated the Ca^2+^-handling parameters in male/female and nSR/AF models to describe the reported sex difference and AF-remodeling of Ca^2+^-handling proteins expression and phosphorylation.^30^ The modifications are summarized in **Table 1** and include the maximum unitary flux and inactivation of L-type Ca^2+^-channel (LTCC, with parameters tuned based on simulated voltage clamp to recapitulate the LTCC changes by AF in male cardiomyocytes^30^), the expression of CSQ, phospholamban (PLB), and RyR2 phosphorylation (**Table 1**). Corresponding parameter values are given in **Supplementary Table S1**.

### Simulation protocol and analysis of Ca^2+^ sparks and SCRs

To investigate the rate dependence of Ca^2+^ dynamics and SCR events following burst pacing, we employed an AP clamp protocol. The protocol consisted of a 4-s unstimulated period, followed by a 28-s AP-clamp period, during which the cardiomyocyte was paced at 0.5, 1, 2, 3, 4, or 5 Hz using AP-clamp protocols. The protocol concluded with a 5-s non-stimulation pause period to allow for evaluation of SCR events. The AP traces used in the protocol at each pacing rate were obtained from current-clamp simulations using the Zhang et. al.^36^ model. The holding potential during the non-stimulated period was set to the resting membrane potential of the AP clamp trace.

To identify SCRs, we analyzed the time course of the averaged [Ca^2+^]_Cyto_ of each line scan, from the beginning of the last beat to the end of the 5^th^ second no-stimulation period, then excluded the first [Ca^2+^]_Cyto_ peak (i.e., AP-induced Ca^2+^ release). The incidence of Ca^2+^ sparks was determined for each CRU on selected line scans by analyzing the time course of CRU-specific [Ca^2+^]_Cyto_ during the 15-s no-stimulation period. In both cases, the respective [Ca^2+^]_Cyto_ time courses, sampled at 1 kHz, were denoised using a moving-averaging filter (window size = 5). To determine the incidence and amplitude of SCRs, and latency of the first SCR, as well as the incidence of Ca^2+^ sparks, we utilized and the ‘findpeak’ function in MATLAB R2021(MathWorks, Natick, MA, USA), with ’MinPeakProminence’ of 0.05 μM and ’MinPeakDistance’ of 50 ms for the findpeak function to define SCRs for the averaged line scan [Ca^2+^]_Cyto_, and Ca^2+^ sparks for each CRU [Ca^2+^]_Cyto_. The amplitudes of SCR events were calculated as the difference between the peak and the minimum [Ca^2+^]_Cyto_ values during the no-stimulation period. To minimize the dependence of detected Ca^2+^ sparks and SCR events on the location of line scan, we monitored the [Ca^2+^]_Cyto_ along 13 transversal lines selected from the middle z-stack of simulated cardiomyocytes, with a longitudinal spacing of 2.7 μm (i.e., 3 CRUs) between the line scans. To reduce computational costs, modeling data are generated from a single cardiomyocyte model simulation for each group and condition. This approach is justified by the fact that simulation results remain unaffected by model stochasticity, as confirmed through repeated simulations with different random number seeds in our previous study.^36^

### Simulation of varying individual Ca^2+^-handling determinants and candidate therapeutic interventions

We investigated the contribution of each sex- and AF-associated change to atrial arrhythmogenic events through sensitivity analysis by varying one factor at-a-time. The targets included cell width, t-tubular density, the relative change of PLB expression, RyR2 phosphorylation, and LTCC activity. In addition, we simulated several interventions targeting atrial Ca^2+^ handling to demonstrate potential sex differences in anti-arrhythmia intervention outcomes. These interventions included t-tubular system restoration, PLB and CSQ overexpression, sarcoplasmic/endoplasmic reticulum Ca²⁺-ATPase (SERCA) inhibition, and RyR2 dephosphorylation. A detailed descriptions of the parameters for these interventions are listed in **Table 1** and values given in **Supplementary Table S2**.

### Numerical methods, statistical analysis, and data availability

Our sex-specific atrial cardiomyocyte models were implemented in C++ and parallelized using OpenMP 5.1 (https://www.openmp.org/), and with a combination of deterministic and stochastic modeling as done previously.^36,37^ We improved the accuracy of stochastic simulation of the Markov model of RyR2s by replacing the binomial-distribution-based random generator (which was applied to simulate the stochastic transition of RyR2s among the 4 RyR2 Markov states) with a multi-nominal stochastic generator, as implemented in Numpy.^42^ The source code of our new 3D human atrial cell model can be accessed from http://elegrandi.wixsite.com/grandilab/downloads and https://github.com/drgrandilab. Statistical analysis was performed by One-way ANOVA with planned comparison and Bonferroni correction using MATLAB (Version 2022b, The MathWorks, Natick, MA, USA) with a threshold significance level of 0.05.

## Results

### Simulated female vs male AF cardiomyocytes exhibit a greater incidence of Ca^2+^ sparks at rest, similar to experimental observation

We constructed male- and female-specific spatially detailed stochastic models of Ca^2+^ handling for both nSR and chronic AF (cAF) conditions within a human atrial cardiomyocyte model. To test the robustness of our sex- and AF-specific parameterization, we first compared the incidence of subcellar Ca^2+^ sparks in the male and female atrial cardiomyocyte models at rest with experimental data from human samples. The membrane voltage of the model cardiomyocytes were clamped to -72 mV, which corresponds to the resting membrane potential of the baseline model^36^ at 0.5 Hz of steady-state pacing. Transversal line scan analyses (**Fig. 1Ai,ii**) of simulated [Ca^2+^]_Cyto_ revealed increased Ca^2+^ spark incidence in AF vs nSR conditions (**Fig. 1B**) in both male and female cardiomyocytes. There was no sex difference in the Ca^2+^ spark frequency in nSR conditions. However, in AF-remodeled cardiomyocytes the female model exhibited substantially greater Ca^2+^ spark abundance compared with the male model. These results are confirmed in additional simulations wherein the cardiomyocyte resting membrane potential was clamped at various levels (**Supplementary Fig. S1**). Importantly, the AF- and sex-dependent differences in Ca^2+^ spark incidence observed in our simulations qualitatively align with a previous experimental study comparing Ca^2+^ sparks in resting atrial cardiomyocytes from male and female patients with and without AF (**Fig. 1C**).^30^ These simulations suggest that the sex- and AF-related differences incorporated in our models recapitulate the sex differences observed in atrial Ca^2+^ spark properties.

**Fig. 1.**
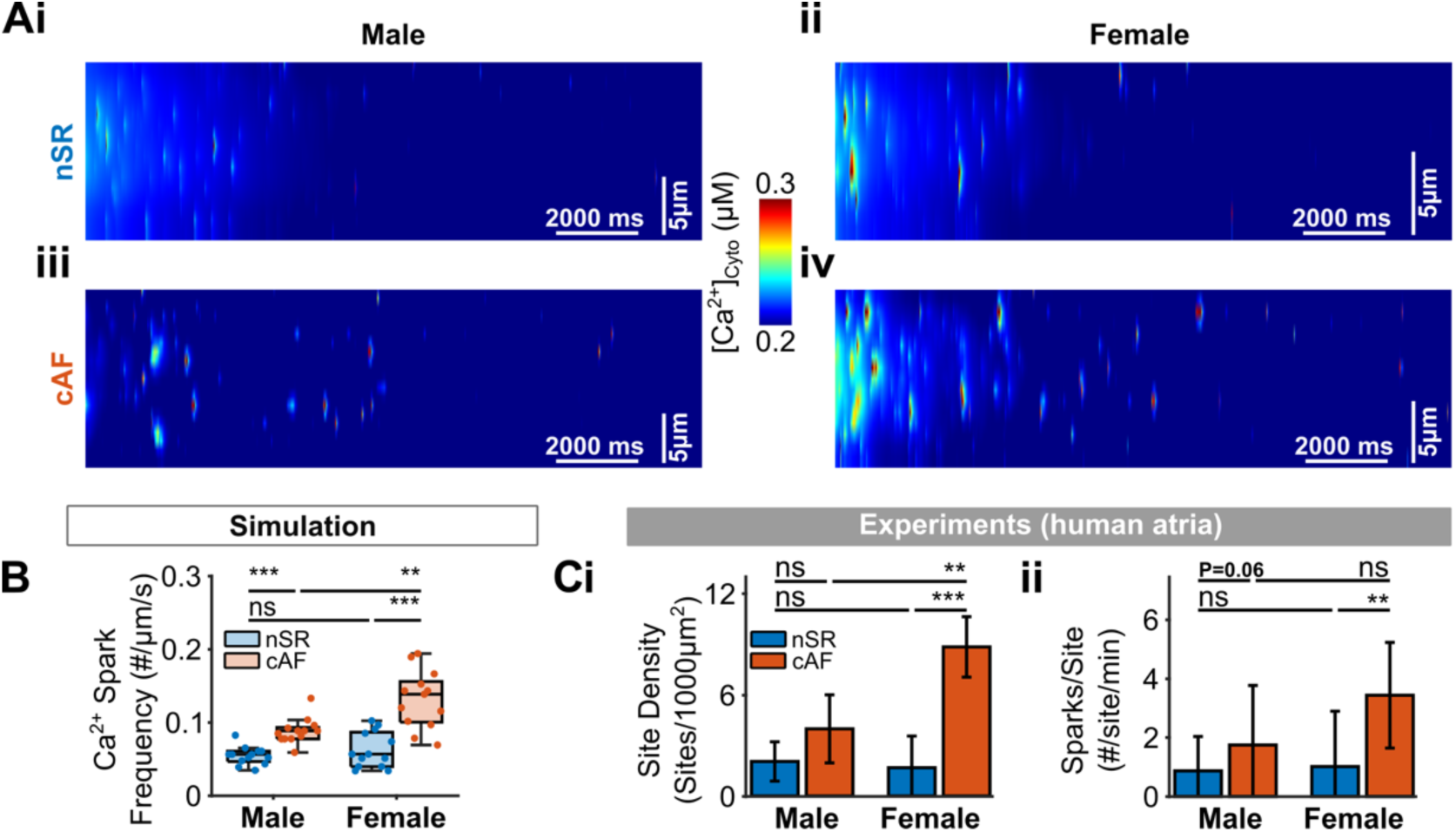
Simulated effects of sex and AF on Ca^2+^ sparks and comparison to experiments. (**A**) Simulated transversal line scan of intracellular Ca^2+^ from **(i, iii)** male and (**ii, iv**) female atrial cardiomyocytes in (**i, ii**) nSR and (**iii, iv**) cAF conditions. (**B**) Quantification of simulated Ca^2+^ spark frequency. For each group, data are reported from the analysis of 13 line scans in 1 cardiomyocyte model simulation. Statistical significance was determined by One-way ANOVA with planned comparison and Bonferroni correction and indicated with ***: p < 0.001; **: p < 0.01; and ns: not significant. (**C**) Ca^2+^ spark (**i**) site density and (**ii**) number per site from human atrial cardiomyocyte experiments reported by Herraiz-Martinez et al.^30^.

### Fast atrial rate-induced SCR events are more frequent in female vs male cardiomyocytes in AF

The elevated Ca^2+^-spark incidence in female AF may be indicative of arrhythmogenic Ca^2+^-driven instabilities in female vs male atria. However, although sex difference in Ca^2+^ sparks has been reported from quiescent human atrial cardiomyocytes,^30^ SCRs and their rate-dependence that is critically relevant to AF, have not yet been investigated. As such, we next sought to determine the arrhythmogenic propensity of atrial cardiomyocytes using a pacing-pause protocol. Because experimental characterization of sex differences in atrial electrophysiology is lacking, and a recent study reported no difference between male and female AP characteristics in AF at 1 Hz,^43^ we applied an AP-clamp protocol to provoke SCRs using the same AP configuration for both sexes.

This allowed studying the effects of sex differences in Ca^2+^ handling independent of possible sex differences in atrial electrophysiology. To achieve this, we generated trains of APs by pacing our baseline cardiomyocyte model^36,37^ at various rates and applied these APs to all cardiomyocyte models prior to holding at the resting membrane potential to record SCRs (**Fig. 2Ai**). Transversal line scans of [Ca^2+^]_Cyto_ were obtained (**Fig. 2Aiii-iv**) to assess local SCRs, which were identified as post-repolarization instability in the averaged line-scan [Ca^2+^]_Cyto_ (**Fig. 2Av**). We found that the stimulation protocol led to SCRs in both male and female atrial cardiomyocytes (**Fig. 2Aiii,iv**). There was no sex-dependent difference in SCR incidence in nSR conditions except for the 5-Hz AP clamp (**Fig. 2B**). However, in AF conditions female cardiomyocytes exhibited a greater incidence of SCRs compared to male cardiomyocytes (**Fig. 2Aiv,v** and **2Bi-iv**). Of note, the sex-dependent difference in SCR incidence was not attributed to differences in SR load, which was similar between male and female cardiomyocytes in nSR, and slightly lower in the female vs male cardiomyocytes in AF (**Supplementary Fig. S4Biv**). Interestingly, AF caused a reduction (vs nSR) in the maximum amplitude of SCRs in male cardiomyocytes (**Fig. 2Ci-iv**), owing to enhanced SERCA activity (due to PLB reduction) and whole-cell Ca^2+^ unloading due to LTCC reduction (see **Graphical abstract**). In contrast, the maximum SCR amplitude was elevated in AF-vs-nSR female cardiomyocytes (**Fig. 2Ci-iv**), due to hyperactive RyR2 (see **Graphical abstract**). Overall, the sex-dependent differences in AF were amplified with faster AP rates, displaying a positive rate dependence (**Fig. 2Bi-iv,Ci-iv**). We also found that in male AF the latency of SCRs was delayed compared to female cardiomyocytes and nSR conditions (**Fig. 2Di-iv**). Collectively, these simulations showed sex differences in the incidence of potentially proarrhythmic SCRs in human AF and pointing to a higher susceptibility to Ca^2+^-dependent arrhythmias in female vs male cardiomyocytes.

**Fig. 2.**
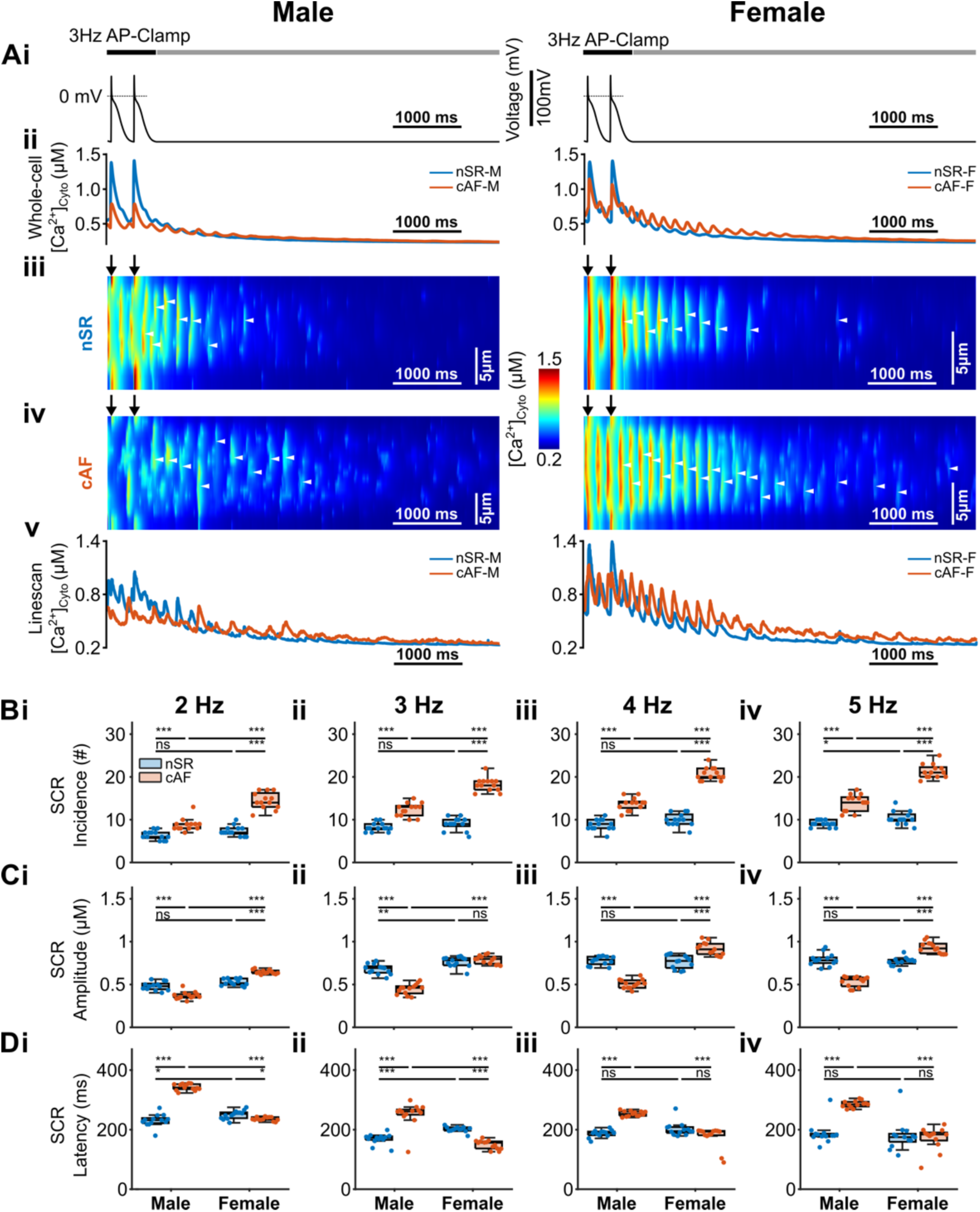
Simulated effects of sex and AF on spontaneous Ca^2+^ release (SCR) events in human atrial cardiomyocytes. **(A**) **(i)** AP-clamp traces at 3 Hz and subsequent resting period, **(ii)** whole-cell averaged cytosolic Ca^2+^, **(iii-v)** transverse line scan of cytosolic Ca^2+^ in **(iii)** nSR vs **(iv)** cAF, and **(v)** the corresponding averaged line-scan Ca^2+^ traces for male (*left*) vs female (*right*) cardiomyocytes. White arrow heads indicate SCRs and black arrows mark the start of an AP. (**B-C**) Summary of (**B**) SCR incidence, (**C**) SCR amplitude, and (**D**) Latency to the first SCR in female vs male cardiomyocytes in AF conditions. Columns (i-iv) of (**B-D**) depict data from AP clamps of 2 Hz to 5 Hz, respectively. For each group in (**B-D**), data are reported from the analysis of 13 line scans in 1 cardiomyocyte model simulation. Statistical significance was determined by One-way ANOVA with planned comparison and Bonferroni correction and indicated with ***: p < 0.001; **: p < 0.01; *: p < 0.05; ns: not significant.

### Sensitivity analysis unmasks the individual contribution of sex- and AF-dependent differences to SCR

Our simulations revealed sex-dependent differences in propensity to SCRs due to the combined effects of sex-related and AF-induced differences in subcellular Ca^2+^-handling. To dissect the contribution of individual alterations, we performed parameter sensitivity analysis by varying one parameter at-a-time in the nSR male-specific model (**Fig. 3**) under AP trains (3-Hz pacing) applied to provoke SCRs. Simulating varying cell width did not affect SCR incidence, with a trend of augmented SCR amplitude in wider myocytes (**Fig. 3Ai-ii**). This suggests that the increased cardiomyocyte size in AF vs nSR conditions or male vs female atria may have a limited contribution to the simulated disease- and sex-dependent phenotypes. Conversely, decreasing t-tubular density substantially increased both incidence and amplitude of SCRs (**Fig. 3Bi-ii)**, indicating a potential proarrhythmic role of the t-tubular loss noted in AF. Indeed, our previous work demonstrated that loss of tubules promoted SCRs in paced cardiomyocytes through reduction of NCX-dependent Ca^2+^ export, causing diastolic Ca^2+^ accumulation and increased RyR2 open probability.^36^ These effects were stronger for the RyR2s located at the cell interior than those at the periphery and enhanced by faster AP rates.^36^ Taken together, these results suggest that the loss of t-tubules promotes arrhythmic events in both sexes.

**Fig. 3.**
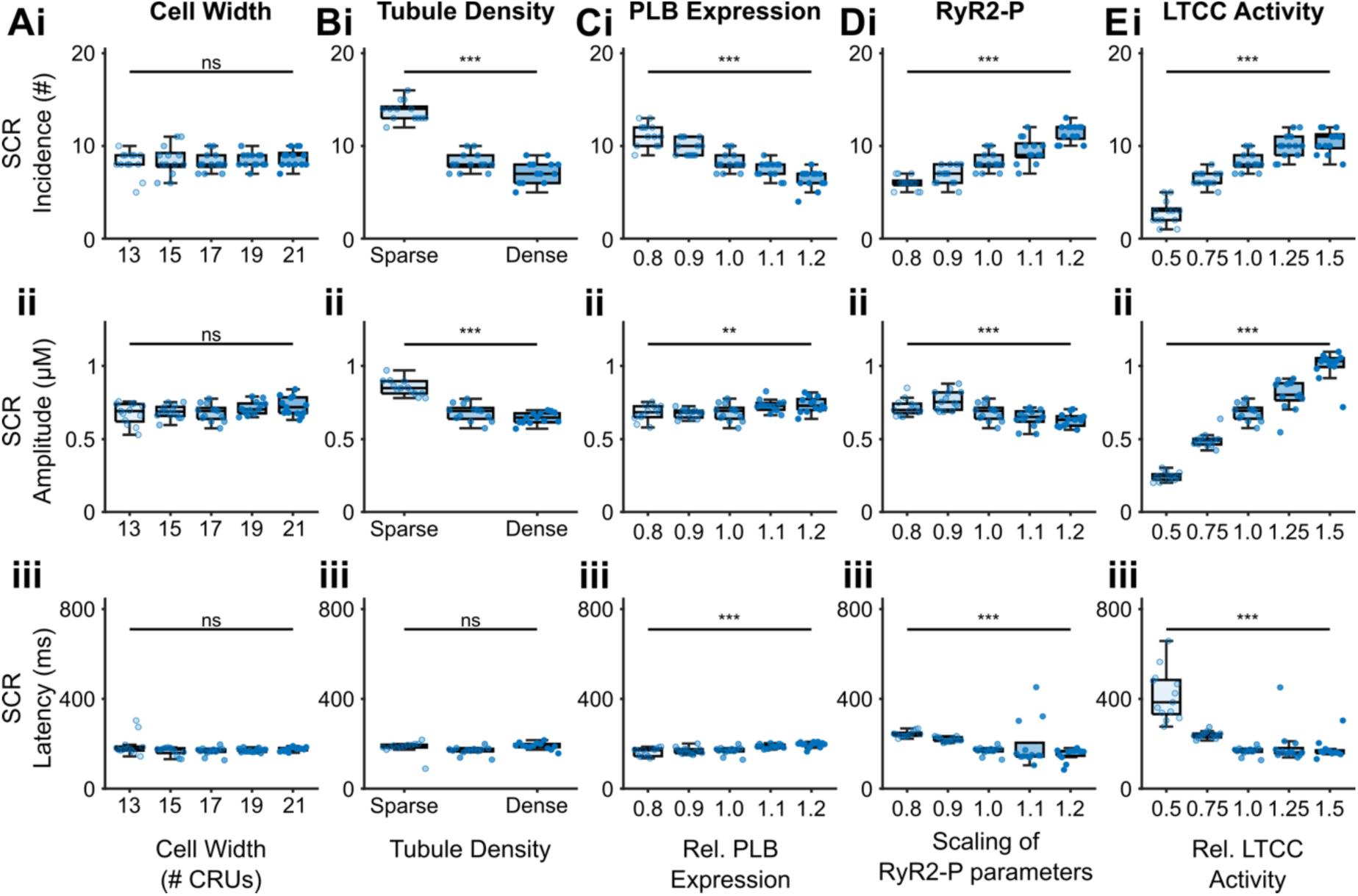
Individual contribution of sex- and AF-dependent alterations to SCR event formation in human atrial cardiomyocytes determined by changing one parameter at-a-time. (**A-E**) Effects of varying **(A)** cell width, **(B)** t-tubule density, **(C)** relative change of PLB expression, **(D)** relative change of RyR2 phosphorylation, and **(E)** relative change of LTCC activity, on **(i)** SCR incidence, **(ii)** SCR amplitude, and **(iii)** SCR latency simulated using male cardiomyocyte model in nSR conditions. For each group, data are reported from the analysis of 13 line scans in 1 cardiomyocyte model simulation. Statistical significance was determined by One-way ANOVA with planned comparison and Bonferroni correction and indicated with ***: p < 0.001; **: p < 0.01; *: p < 0.05; ns: not significant.

Similarly, reducing PLB expression (e.g., as seen in male AF cardiomyocytes) caused moderate elevation of SCR incidence, while slightly decreasing SCR amplitude (by promoting SERCA activity) and latency (**Fig. 3Ci-iii**). Our simulations showed that increased RyR2 phosphorylation (as noted in the female atria^30^) alone moderately increased SCR incidence and shortened SCR latency, although with slightly lower amplitude (**Fig. 3Di-iii**). Furthermore, reducing L-type Ca^2+^ channel activity drastically diminished SCR incidence and amplitude and delayed SCR latency (**Fig. 3Ei-iii**), due to unloading of the cytosolic and SR Ca^2+^ (see **Graphical abstract**). Finally, our previous modeling work demonstrated a SCR-promoting effect of CSQ downregulation.^37^ Therefore, the AF-associated reduction in PLB expression in male atria and elevated RyR2 phosphorylation in the female atria augmented SCRs in male and female cardiomyocytes, respectively, whereas the AF-induced reduction of L-type Ca^2+^ channel activity in the male atria could ameliorate the propensity to SCRs, thus contributing substantially to the female-vs-male increase in Ca^2+^-dependent arrhythmias in AF.

### Interventions targeting individual Ca^2+^-handling determinants alleviates SCRs

Correcting dysfunctional intracellular Ca^2+^ signaling in AF is a promising anti-AF strategy.^36–39^ As our simulations predicted the increase in AF-associated arrhythmogenic SCRs in female vs male atrial cardiomyocytes, we next sought to evaluate their responses to putative Ca^2+^-targeted interventions. We simulated the upregulation of t-tubular structure (Tubule up), CSQ (CSQ up) and PLB expression (PLB up), and downregulation of RyR2 (RyR2 down, i.e., dephosphorylation) and SERCA (SERCA down, i.e., inhibition) (**Fig. 4**). As above, we applied a 3 Hz AP-clamp protocol (**Fig. 4A,B**) to compare the SCR characteristics in AF cardiomyocytes with the proposed interventions. We found that all putative interventions could suppress SCR incidence (**Fig. 4C**) and delay SCR latency (**Fig. 4E**), with t-tubule recovery and CSQ upregulation also decreasing SCR amplitude in female cardiomyocytes (**Fig. 4Di,iv**). We define the effectiveness of an intervention as a reduction of SCR incidence post-intervention reaching a level comparable to or below the baseline observed in male cardiomyocytes under nSR conditions (shown with dashed lines in **Fig. 4C-E**). T-tubule recovery (**Fig. 4Bi, Ci**), RyR2 dephosphorylation (**Fig. 4Cii, *Supplementary Fig. S2B***), and SERCA inhibition (**Fig. 4Ciii, *Supplementary Fig. S2C***) alone were effective in preventing SCRs in both sexes, with greater SCR-suppressing responses in female vs male cardiomyocytes. In contrast, PLB overexpression was effective in male but not female cardiomyocytes (**Fig. 4Bii, Cv**). CSQ overexpression alone was not effective against SCR in either sex (***Supplementary Fig. S2D,* Fig. 4Civ**).

**Fig. 4.**
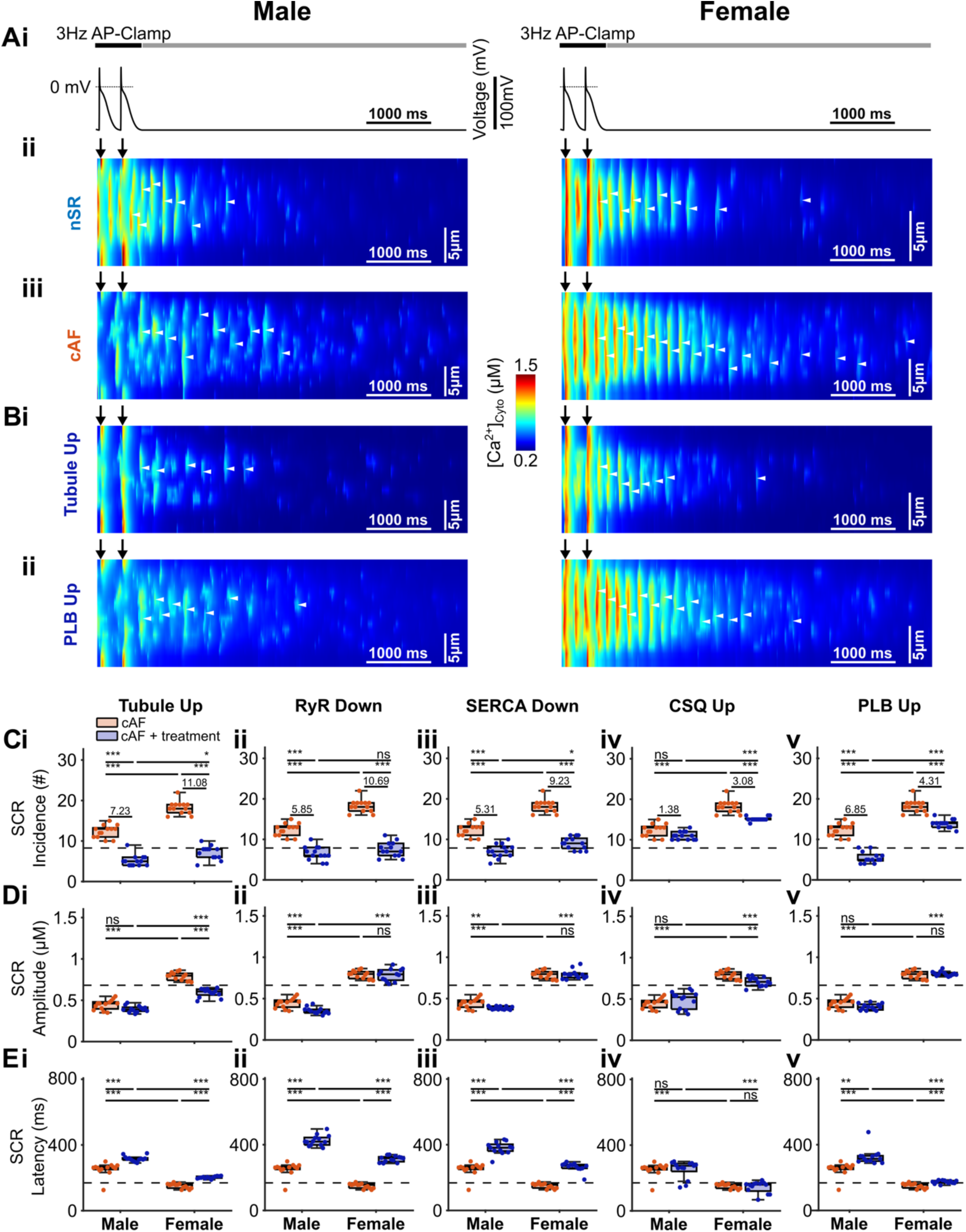
Simulated effects of targeted interventions of Ca^2+^-handling determinants in male and female atrial cardiomyocytes from AF patients. **(A) (i)** AP-clamp traces and subsequent resting period, **(ii-iii)** simulated transversal line scans of [Ca^2+^]_Cyto_ for male and female cardiomyocytes in **(ii)** nSR and **(iii)** AF conditions as in Fig. 2A are illustrated for comparison. **(B)** Simulated transversal line scans of [Ca^2+^]_Cyto_ with interventions upregulating (**i**) t-tubular density and (**ii**) PLB in human atrial cardiomyocytes with AF. White arrowheads indicate SCRs and black arrows mark the start of AP. **(C-E)** Summary of intervention effects on (**C**) SCR incidence, (**D**) SCR amplitude, and (**E**) SCR latency; columns (**i-v**) illustrate effects of proposed interventions with (**i**) Tubule up, (**ii**) RyR2 down, (**iii**) SERCA down, (**iv**) CSQ up, and (**v**) PLB up. In Panel (**C**), the numbers indicate the mean reduction in SCR incidence due to intervention. For each group in (**C-E**), data are reported from the analysis of 13 line scans in 1 cardiomyocyte model simulation. Statistical significance was determined by One-way ANOVA with planned comparison and Bonferroni correction and indicated with ***: p < 0.001; **: p < 0.01; *: p < 0.05; ns: not significant.

Next, we performed additional simulations to test the effects of multi-target interventions with pairwise combination of the proposed maneuvers (**Fig. 5A-E** and **Supplementary Fig. S3A-E**). In general, compared with the single-hit interventions (**Fig. 4**), combined interventions caused a stronger SCR suppression (**Fig. 5C**), led to greater reductions of SCR amplitude (to values lower than nSR baseline, **Fig. 5D**), and stronger prolongation of SCR latency (**Fig. 5E**). Overall, the extent of SCR suppression was more pronounced in female vs male cardiomyocytes. Interestingly, CSQ overexpression presented an exception, whereby the combined interventions of CSQ overexpression with another intervention resulted in less effective outcomes than with the standalone application of the individual intervention (**Supplementary Fig. S3Cii-v**). Collectively, our simulations suggest that the combined interventions that correct t-tubular structure, RyR2 phosphorylation, SERCA activity and PLB expression, are more effective in suppressing SCRs in compared to a single-hit approach, with female cardiomyocytes showing additive responses to these combined interventions.

**Fig. 5.**
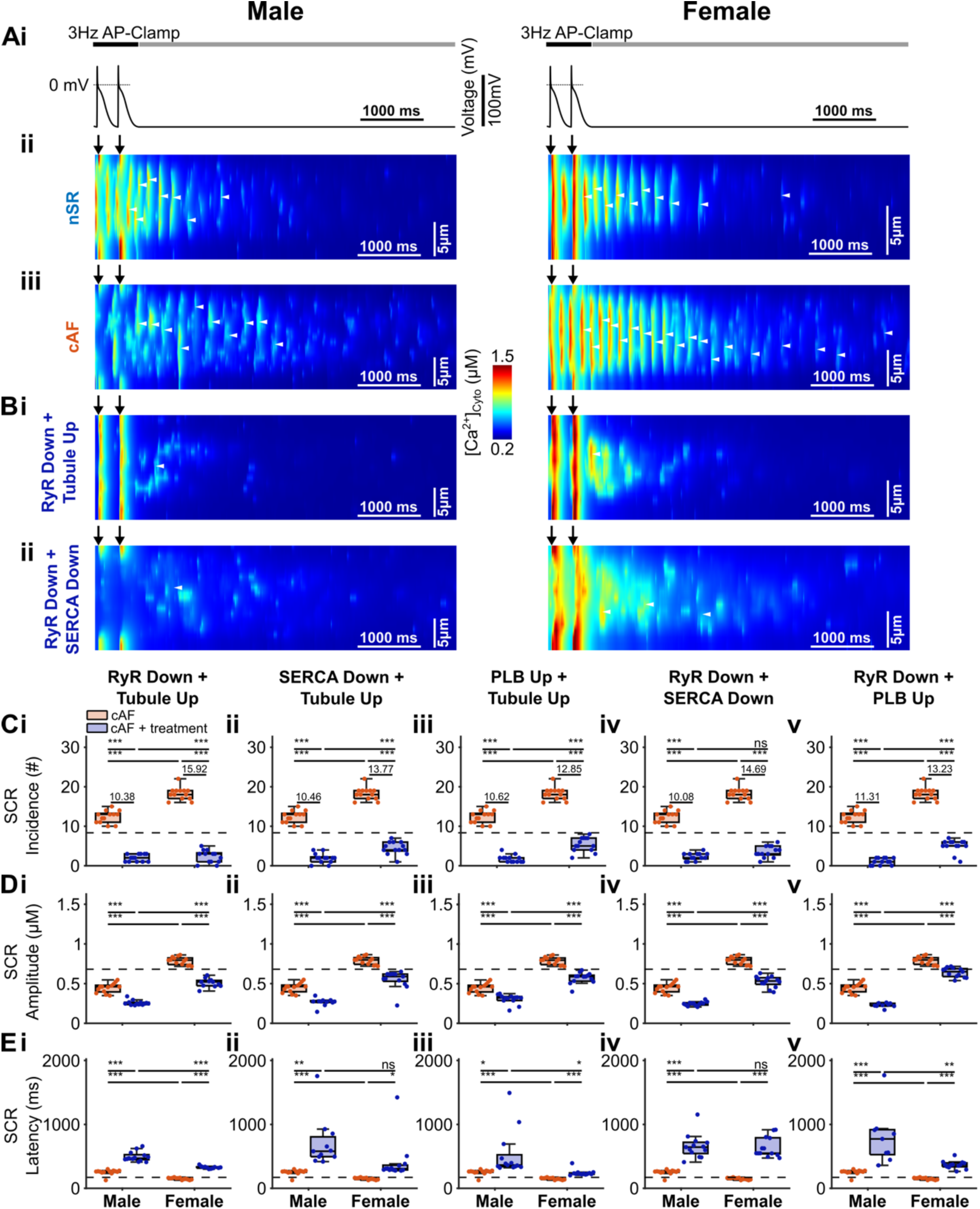
Simulated effects of pair-wise combined interventions on male and female atrial cardiomyocytes from AF patients. **(A) (i)** AP-clamp traces, **(ii-iii)** simulated transversal line-scan [Ca^2+^]_Cyto_ for male and female cardiomyocytes in **(ii)** nSR and **(iii)** AF conditions as in Fig. 2A are illustrated for comparison. **(B)** Simulated transversal line scans of [Ca^2+^]_Cyto_ with combined interventions of (**i**) RyR2 down + Tubule up, and (**ii**) RyR2 down + SERCA down. White arrow heads indicate SCRs and black arrows mark the start of AP. **(C-E)** Summary of intervention effects on **(C)** SCR incidence, **(D)** SCR amplitude, and **(E)** SCR latency; columns **(i-v)** depict effects of combined intervention with (**i**) RyR2 down + Tubule up, (**ii**) SERCA down + Tubule up, (**iii**) PLB up + Tubule up, (**iv**) RyR2 down + SERCA down, and (**v**) RyR2 down + PLB up. In Panel (**C**), the numbers indicate the mean reduction of SCR incidence due to intervention. For each group in (**C-E**), data are reported from the analysis of 13 line scans in 1 cardiomyocyte model simulation. Statistical significance was determined by One-way ANOVA with planned comparison and Bonferroni correction and indicated with ***: p < 0.001; **: p < 0.01; *: p < 0.05; ns: not significant.

### Beat-to-beat Ca^2+^-transient variability is greater in female vs male cardiomyocytes in AF

Our simulations uncovered an increased propensity to SCRs in female vs male cardiomyocytes in AF and demonstrated SCR-suppressing effects of interventions targeting subcellular Ca^2+^-handling determinants. However, how sex modifies the characteristics of the Ca^2+^ transient (CaT) elicited during APs, especially in AF, is unknown. Therefore, we compared the AP-evoked CaT properties in our male and female atrial cardiomyocyte models in both nSR and AF conditions, by clamping the membrane with AP trains at frequencies ranging from 0.5 to 5 Hz (**Supplementary Fig. S4A**). Analyses of whole-cell CaT demonstrated rate-dependent increases of systolic and diastolic Ca^2+^ levels as well as acceleration of relaxation for the cardiomyocyte models from both sexes (**Supplementary Fig. S4B**). In both male and female, simulations recapitulated the experimentally observed reduction of CaT amplitude in atrial cardiomyocytes of AF patients.^22,23^

However, the reduction of CaT amplitude in male cardiomyocytes was the consequence of the lower systolic [Ca^2+^]_Cyto_, whereas in the female cardiomyocytes this was due to the higher diastolic [Ca^2+^]_Cyto_ (**Supplementary Fig. S4Bi-ii**). Male cardiomyocytes also displayed a slightly greater systolic [Ca^2+^]_Cyto_ than the female cardiomyocytes in nSR conditions, but this pattern was reversed in AF conditions (**Supplementary Fig. S4A-B**). Finally, the simulated SR Ca^2+^ load was not different between male and female cardiomyocytes in nSR conditions and slightly lower in the female vs male cardiomyocytes in AF conditions. These observations in the SR load are consistent with previous experimental measurements using the time-integral of the caffeine-induced (NCX) current.^30^

Beat-to-beat variability of the CaT during pacing cycles may induce AP instabilities that could promote arrhythmogenesis. We sought to determine if the more frequent diastolic SCRs do increase beat-to-beat variability of the atrial CaT. The simulated transversal line scans of subcellular CaT measured at the center of the cell revealed a greater asynchrony (i.e., U-shaped pattern) in centripetal CaT propagation for male vs female cardiomyocytes, a phenomenon that was promoted by AF in both sexes (**Fig. 6Aii-iii**). Like the SCR elicited following pacing-pause AP-clamp, these transversal line scans of cytosolic Ca^2+^ revealed diastolic SCRs between the AP-induced CaTs (**Fig. 6Ai-iv**). The incidence of diastolic SCRs was higher for AF vs nSR conditions in the female but not male cardiomyocytes (**Fig. 6Aii-iv**), consistent with the greater incidence of SCRs after pacing in the female cardiomyocytes in AF (**Fig. 2B**).

**Fig. 6.**
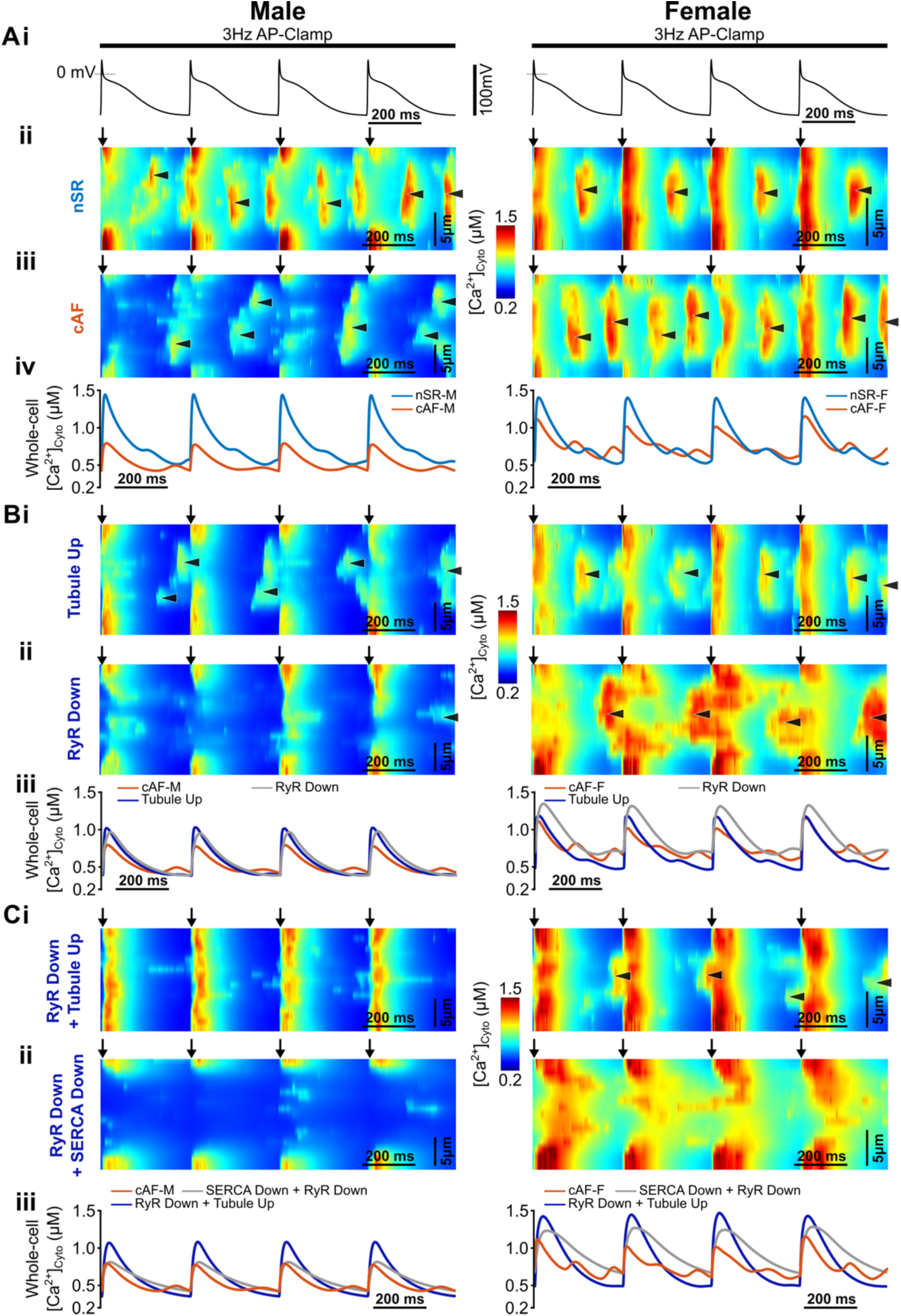
Simulated transversal line scans of AP-clamp evoked Ca^2+^ transients in male and female atrial cardiomyocytes and responses to interventions in AF. **(A) (i)** AP-clamp traces, **(ii-iv)** simulated transversal line scans of AP clamp-elicited CaTs for male and female cardiomyocytes in **(ii)** nSR and **(iii)** AF conditions, and **(iv)** whole-cell averaged CaTs. **(B)** Simulated transversal line scans of AP clamp-elicited CaTs from cardiomyocytes in AF following interventions causing **(i)** Tubule up, **(ii)** RyR2 down, and **(iii)** comparison of whole-cell averaged CaTs with and without intervention. **(C)** Simulated transversal line scans of AP clamp-elicited CaTs cardiomyocytes in AF following combined interventions of **(i)** RyR2 down + Tubule up, **(ii)** RyR2 down + SERCA down, and **(iii)** comparison of whole-cell averaged CaTs with and without intervention. Arrows mark the start of AP, and the diastolic SCRs are indicated with arrow heads. For each group, data are reported from 1 cardiomyocyte model simulation.

We found that the diastolic SCRs show a beat-to-beat variation in incidence and amplitude, especially in AF (**Fig. 6Aii-iii**). This variation may impact SR load and RyR2 refractoriness and could increase the beat-to-beat variability in AP-coupled CaT amplitude. We computed the beat-to-beat variation of whole-cell CaT amplitude evoked using AP clamp at 0.5 to 5 Hz (**Supplementary Fig. S5A-B**). While the beat-to-beat variation in CaT amplitude was similar between the two sexes in nSR conditions, AF simulations unmasked sex-dependent changes. Elevated beat-to-beat CaT variation was observed at fast rates in AF vs nSR in female cardiomyocytes (3 to 5 Hz), whereas beat-to-beat CaT variation was reduced in male cardiomyocytes (4 and 5 Hz). Thus, at fast rates beat-to-beat CaT variability is greater in female vs male AF cardiomyocytes, pointing to a greater Ca^2+^-handling instability in the female atrial cardiomyocytes in AF.

Sensitivity analysis revealed the impact of individual sex-dependent differences on the beat-to-beat CaT variability (**Supplementary Fig. S5C**). We found that female-to-male differences, *i.e.,* lower cell width and t-tubular density, but greater LTCC activity and RyR2 phosphorylation, could all contribute to greater beat-to-beat CaT variability in female vs male cardiomyocytes, whereas the PLB loss alone (as noted in male cardiomyocytes) could elevate the beat-to-beat CaT variability for the male cardiomyocytes (**Supplementary Fig. S5Ciii**).

We also evaluated how the Ca^2+^-targeted interventions impacted the beat-to-beat CaT variability in AF (**Fig. 6B,C, Supplementary Fig. S6**). Simulated line-scan [Ca^2+^]_Cyto_ showed that correcting RyR2 (**Fig. 6Bii**), SERCA, PLB, or CSQ alone (**Supplementary Fig. S6Aiii-v**) did not affect the asynchronized, U-shaped subcellular CaT propagation. In contrast, t-tubule recovery (either increasing t-tubular density alone or combined with an additional intervention) improved the synchronization of AP-evoked subcellular CaTs (**Fig. 6Bi**). In general, the simulated interventions reduced the incidence of diastolic SCRs in both sexes (**Fig. 6B,C, Supplementary Fig. S6Aiv-viii**); however, upregulation of CSQ instead promoted diastolic SCRs in female cardiomyocytes (**Supplementary Fig. S6Aiii**). The efficacy to suppress diastolic SCRs was enhanced with combined compared to single-hit interventions, especially for interventions that involved t-tubular recovery in the male cardiomyocytes (**Fig. 6C, Supplementary Fig. S6Avi-viii**).

**Fig. 7** and **Supplementary Fig. S7** summarize the effects of these interventions on the AP-induced CaT amplitude, CaT beat-to-beat variability, and the post-AP SCR incidence, in female and male atrial cardiomyocytes with AF, respectively. Indeed, these interventions generally could suppress SCR incidence (**Fig. 7A**), and lower SCR amplitude (**Fig. 7B**), increase (**Fig. 7C**) the CaT amplitude, and reduce the beat-to-beat CaT variability (**Fig. 7D**) in female cardiomyocytes. Similar effects were observed in male cardiomyocytes (**Supplementary Fig. S7A-D**). To compare the effects of the tested interventions for female atrial cardiomyocytes in AF, we performed a multi-objective assessment of combined intervention regimens. Given that the male cardiomyocytes displayed a lower incidence of SCRs, we used the phenotype of male atrial cardiomyocytes in nSR as reference to assess the intervention outcomes of the female atrial cardiomyocytes. When solely considering the SCR incidence and amplitude, 8 intervention regimens (out of 15) were deemed effective for female cardiomyocytes (**Fig. 7 A,B**). This number dropped to 5 for female cardiomyocytes when a reduction of beat-to-beat CaT variability was included in the objectives (**Fig. 7D**). The top 6 performing interventions for female atrial cardiomyocytes in AF are ‘double-hit’ interventions involving combinations of t-tubular system improvement with either downregulation of RyR2 or SERCA, or upregulation of PLB (Group 1, **Fig. 7**). Overall, these combined intervention regimens outperformed the respective single-hit interventions. Interestingly, combinations that added CSQ worsened the outcome compared to single-hit interventions without CSQ (Group 2, **Fig. 7**), while sole PLB and CSQ modulations were not effective in female cardiomyocytes (Group 3, **Fig. 7**), and CSQ modulation alone failed also to affect male cardiomyocytes (**Supplementary Fig. S7**). Of note, while all interventions could increase the CaT amplitude (and thus improve the related atrial contractility), interventions that incorporated t-tubular system restoration were generally most effective at elevating CaT amplitude (and thus contractility). Taken together, these results suggest that combining t-tubular structure restoration with correction of Ca^2+^-handling protein abnormalities may be a viable approach to target Ca^2+^-driven arrhythmias in females with AF.

**Fig 7.**
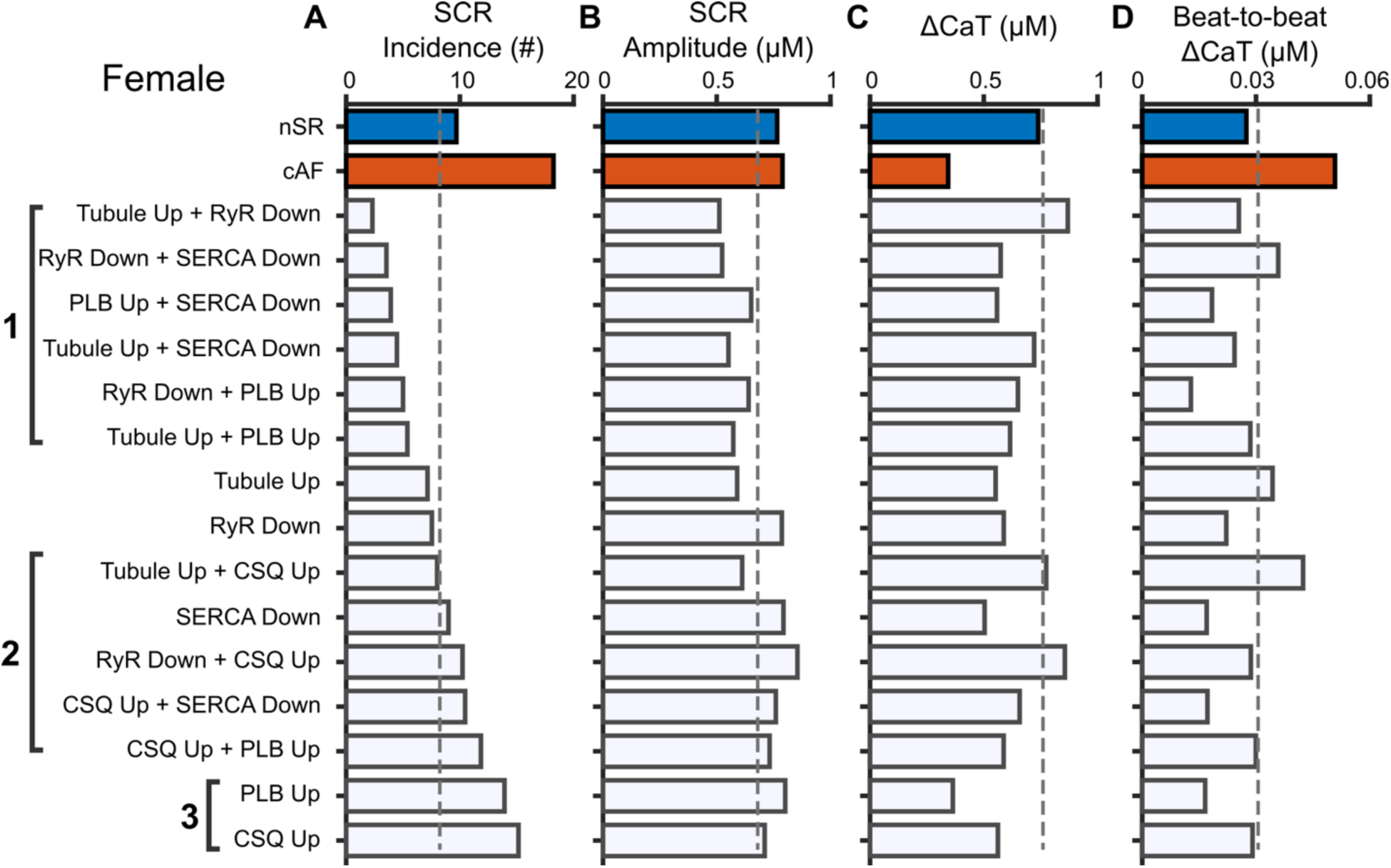
Summary data comparing the intervention effects on SCR (A) incidence and (B) amplitude, (C) whole-cell CaT amplitude, and (D) CaT beat-to-beat variability in female atrial cardiomyocytes. Dashed lines indicate the referencing values obtained from male cardiomyocytes in nSR conditions. Groups 1 – 3 highlight (1) combined interventions targeting t-tubules, RyR2s, SERCA, and PLB, (2) combinations of CSQ intervention with interventions targeting t-tubules, RyR2s, SERCA, or PLB, and (3) interventions with PLB and CSQ overexpression.

## Discussion

A deeper mechanistic understanding of the sex-dependent differences in AF is urgently required to advance the development of tailored, sex-specific approaches to treat this prevalent condition. In this study, we have built a suite of sex-specific models coupling the spatiotemporal dynamics between electrophysiology and subcellar Ca^2+^ signaling. We leveraged our recently developed 3D human atrial cardiomyocyte model to incorporate known sex- and AF-related differences in Ca^2+^ handling and cellular ultrastructure. Our simulations revealed that, in AF conditions, female vs male atrial cardiomyocytes exhibited greater frequency of Ca^2+^ sparks and a higher susceptibility to pacing-induced SCRs. The beat-to-beat variability in the AP-evoked CaT was also greater in female vs male cardiomyocytes. Combined, our simulation data demonstrated a higher propensity to Ca^2+^-driven arrhythmogenic events in female compared to male atrial cardiomyocytes in AF conditions, which was attributed to sex-dependent differences in AF-remodeled components, primarily increased RyR2 phosphorylation in female and reduced LTCC activity in male atrial cardiomyocytes. Furthermore, our modeling identified Ca^2+^-targeted interventions that could attenuate the Ca^2+^-driven arrhythmic events and informed about combined interventions that correct Ca^2+^-handling abnormalities (e.g., RyR2 dephosphorylation, SERCA inhibition, and PLB overexpression) and restore t-tubule system as potentially effective strategies for suppressing Ca^2+^-driven arrhythmias in female patients with AF. Collectively, our study establishes that AF treatment may benefit from sex-dependent therapeutic approaches informed by sex-specific mechanisms.

### Female atrial cardiomyocytes reveal greater propensity to proarrhythmic SCRs

Despite substantial sex-dependent differences in the clinical symptoms, prescribed treatment options, and outcomes of AF, mechanistic studies to uncover the sex-specific underpinnings of AF pathophysiology have been insufficient. Our sex-specific models show greater incidence of Ca^2+^ sparks in female vs male atrial cardiomyocytes with AF at rest (**Fig. 1**), in agreement with previous experimental reports.^30^ Using a pacing protocol that is relevant to studying arrhythmia, our simulations demonstrated that female vs male cardiomyocytes exhibited much greater incidence and amplitude of pacing-induced SCRs in AF conditions, along with a shorter SCR latency (**Fig. 2**), all reflecting more aberrant RyR2 function and pointing to greater propensity to developing Ca^2+^-dependent triggered activities in females. Interestingly, previous clinical observations documented a greater incidence of non-pulmonary vein triggers in female vs male atria with AF.^7^ This may be, at least in part, due to the greater propensity to cellular triggered activity in female vs males that we identified here. Thus, our simulations offer plausible mechanistic insights into this sex-dependent difference in clinical AF presentation. Additionally, our data show that the beat-to-beat variability of AP-elicited CaTs was much greater in female vs male cardiomyocytes in AF, despite being comparable in nSR conditions (**Fig. 6** and **Supplementary Fig. S5)**. The elevated beat-to-beat variability in female cardiomyocytes was paralleled by a greater incidence of diastolic SCRs (**Fig. 6**). Through Ca^2+^/V_m_ bidirectional coupling, the instability in CaT could translate into beat-to-beat variability in APs, and subsequent increase in spatial AP heterogeneity in tissue that promotes vulnerability to AF-maintaining reentry. Combined, our simulation data point to elevated propensity to developing both ectopic activity (through a Ca^2+^-dependent trigger) and reentry (due to beat-to-beat variability in CaT) in female vs male atria in AF. On the other hand, our simulations suggest that the triggered activity mechanism may be less prominent in male atria with AF and imply that reentry (which is caused by LTCC reduction-mediated shortening of AP and refractoriness) may be the dominant AF-promoting mechanism in male patients.

### Sex-specific molecular and ultrastructural signatures cause SCRs in female atria with AF

Sensitivity analyses of the sex-differential components revealed the individual contribution of each element to the greater propensity to Ca^2+^-dependent arrhythmia in female vs male cardiomyocytes. We found that the greater RyR2 phosphorylation in female and reduced LTCC function in male cardiomyocytes are major determinants of the observed sex-dependent differences (**Fig. 3**), positioning aberrant RyR2s as potential targets for AF treatment in female patients. These results are consistent with our previous study suggesting that reported LTCC reduction by AF is an adaptive mechanism of the atria to limit triggered activity, whereas enhanced RyR2 phosphorylation constitutes a maladaptation that drives the arrhythmia.^28^ In this regard, our suite of sex-specific human atrial cardiomyocyte models is unique and instrumental to gain mechanistic insights into the sex-dependent differences in Ca^2+^-driven arrhythmogenesis in human atria, providing a basis to inform putative treatment strategies to correct the deranged Ca^2+^ signals, especially for the female atria.

### Interventions targeting t-tubules and Ca^2+^ handling effectively suppress SCRs in females

Because Ca^2+^-dependent arrhythmogenesis may be the dominant AF-promoting mechanism in female atria, suppression of Ca^2+^-dependent arrhythmia should be a prioritized objective for therapeutic selection and drug development for female patients. As the Ca^2+^-dependent trigger is not a primary target of current AF therapies, the elevated propensity to this trigger may be a reason why female patients respond more poorly to existing treatment options. For example, clinical evidence suggests that AF recurrence rates after ablation were greater in female vs male patients,^44^ which could be potentially explained by the greater incidence of triggered activity in the female atria. Here, our simulated interventions targeting the t-tubular system and the Ca^2+^-handling machinery (**Figs. 5 – 7**) show that suppression of SCR incidence, reduction of beat-to-beat CaT variability, and promotion of subcellular CaT synchronization could constitute promising anti-AF approaches in female patients. The first two would attenuate Ca^2+^ dependent arrhythmia to achieve rhythm control, while the latter could help rescue atrial contractile dysfunction, which is substantially diminished in AF and favors thrombosis. Interestingly, simulated interventions solely targeting PLB, CSQ, or SERCA did not fully correct SCRs, whereas correcting RyR2s or the t-tubule system alone did not rescue the reduced CaT amplitude (**Fig. 7**). Overall, our simulations suggest that combined interventions of multiple therapeutic targets are more effective than single-hit strategies. Interventions combining t-tubular structure recovery, PLB overexpression, and a reduction of SERCA and RyR2 activities, as highlighted in Group 1 in **Fig. 7**, are the most effective strategies to suppress proarrhythmic SCRs, while also increasing CaT amplitude and thus contractility in female atria. Of note, combinations involving CSQ emerged as exceptions, underscoring the complexity of SCR regulation by CSQ due to the dual effects of CSQ on RyR2 gating and luminal Ca^2+^ buffering in the SR.^37^

The Ca^2+^-targeted intervention strategies identified by our modeling work warrant preclinical experimental assessments. For instance, RyR2 inhibition can be achieved by repurposing dantrolene, a clinical drug used as a RyR1 inhibitor, or using *ent-verticilide*, which is a selective RyR2 inhibitor that suppressed AF susceptibility caused by hyperactive RyR2s.^45^ Indeed, dantrolene has been shown to reduce SCRs in human atria and triggered activity in human ventricles.^46–48^ Also, propafenone^49^ and flecainide are found to act through inhibition of RyR2s when treating catecholaminergic polymorphic ventricular tachycardia.^49,50^ Interestingly, propafenone has greater efficacy of pharmacological cardioversion in female patients,^51^ while the efficacy of preventing AF recurrence by class Ic antiarrhythmic drugs (which includes propafenone and flecainide) was greater in female vs male patients after receiving catheter ablation.^52^ In addition, SERCA inhibition may leverage investigational compounds, such as thapsigargin and cyclopiazonic acid.^53^ Although SERCA upregulation has been pursued for treatment of other heart conditions, e.g., heart failure,^54^ here we found SERCA inhibition instead could reduce SR content to attenuate RyR2 activity (resulting in reduced SCRs, **Fig. 4**), and augmented CaT amplitude (**Fig. 7**). Moreover, restoration of t-tubular structure may take advantage of upregulating several proteins that have been implicated in the regulation of t-tubules, namely junctophilin 2 (JPH2),^55^ amphiphysin II (AMPII, also known as bridging integrator 1/BIN-1),^56,57^ and myotubularin (MTM1),^58^ among others. Gene therapies to increase the protein expression have shown improved outcomes in heart failure by augmenting JPH2,^55^ and AAV9 transduction of BIN1 restores t-tubule microdomains to improve cardiac function in stressed^57^ and db/db diabetic mice.^59^ Interestingly, treatment using tadalafil, a PDE5 inhibitor, restored t-tubules by upregulating BIN1 expression.^60^ While these studies primarily focused on the ventricles, similar treatments and effects may be translated into the atria. Lastly, although reports have been limited, gene therapy may also be applied to augment PLB expression. As such, the therapeutic strategies identified by our modeling work are all experimentally tractable in the future.

### Inclusion of female subjects is key to define sex-specific disease mechanisms

Accumulating evidence show a much lower representation of female sex in both basic and clinical research of cardiovascular diseases,^61,62^ which naturally leads to an inappropriate extrapolation of study outcomes from men to women.^62^ Therefore, a more inclusive representation of female subjects in research is imperative to identify sex-specific disease mechanisms and facilitate the development of targeted treatment strategies considering the unique physiological and pathological aspects of cardiovascular health in female patients. Aligned with this theme, our study reveals increased Ca^2+^-driven arrhythmia mechanisms in female patients with AF, and suggests specific correction of aberrant Ca^2+^-dependent processes as a potential novel therapeutic intervention. Moreover, our study provides novel sex-specific computational models as powerful integrative tools to study emerging biological variables and (patho)physiological mechanisms that are dependent on sex.

### Study limitations

Our modeling approach has several limitations that warrant future investigations. First, our assessment utilized sex-independent AP clamp to reveal Ca^2+^-driven arrhythmic events in the absence of well-established knowledge of sex-dependent differences in atrial APs. Interestingly, recent characterization of atrial APs from male and female patients revealed some sex-dependent differences in AP duration and resting membrane potential; however, these differences were only present in nSR conditions and were diminished in AF.^43^ Additionally, previous expression profiling of human hearts identified several ion channels that were sex-differentially expressed or modified.^30,63^ Therefore, future studies should integrate additional sex-dependent differences in electrophysiology to extend our sex-specific models to investigate the sex-specific interactive role of AP and Ca^2+^ processes in promoting AF and as therapeutic targets. Second, our current sex-specific modeling framework incorporates the direct effectors of electrophysiology and Ca^2+^ signaling. As previous studies involving animals uncovered sex-dependent differences in upstream signals including cAMP/phosphodiesterase in the ventricles,^64,65^ similar differences, if confirmed in the atria, can be readily incorporated into our framework, as established in our atrial signaling model.^28^ Third, our current work solely focused on isolated cardiomyocytes. Previous studies associated female atria with greater fibrosis^44,66^ as compared to male atria. Thus, the increased incidence of Ca^2+^-dependent triggers in female atria as revealed by our study may be exacerbated in tissue through mismatched source-sink relation to trigger spontaneous APs and/or subthreshold delayed afterdepolarizations. The latter could promote tissue dispersion of cell excitability due to heterogeneous availability in fast Na^+^ current due to heterogeneous voltage and Ca^2+^-dependent signaling,^28^ thereby causing functional conduction block that promotes reentry.^67^ The implications for spontaneous APs and/or subthreshold delayed afterdepolarizations warrant future investigations using sex-specific tissue models of human atria.

### Conclusions

In summary, we have built a family of male- and female-specific 3D models of human atrial cardiomyocytes in both nSR and AF conditions. Our analyses demonstrated increased propensity to developing Ca^2+^-driven arrhythmias in female vs male atrial cardiomyocytes with AF, and suggested Ca^2+^-targeted interventions as promising strategies for female AF patients. Our study advocates for more inclusion of female subjects and provides new sex-specific computational models as useful means to define sex-specific disease mechanisms and inform the development of sex-specific therapeutics, ultimately improving effectiveness and equity of AF management.

## Funding and Acknowledgements

### Funding

This work was supported by American Heart Association Postdoctoral Fellowships 20POST35120462 (H.N.), 24POST1195229 (Y.W.), and Predoctoral Fellowship 20PRE35120465 (X.Z.); National Institutes of Health/NHLBI Grants R01HL131517 (E.G., W.E.L., and D.D.), R01HL170521 (E.G., H.N.), R01HL141214 and P01HL141084 (E.G.), R01HL136389 (D.D.), R01HL089598 (D.D.), R01HL163277 (D.D.), R01HL160992 (D.D.), R01HL165704 (D.D.), R00HL138160 (S.M.); NIH Stimulating Peripheral Activity to Relieve Conditions Grant 1OT2OD026580-01 (E.G.); the Research Council of Norway (Grant No. 287395, W.E.L.); UC Davis School of Medicine Dean’s Fellow Award (E.G.); the European Union (large-scale integrative project MAESTRIA, No. 965286, D.D.); Burroughs Wellcome Fund - Doris Duke Charitable Foundation “COVID-19 Fund to Retain Clinical Scientists” Award (S.M.).

