## Supplemental Materials for "Enhanced Ca^2+^-Driven Arrhythmias in Female Patients with Atrial Fibrillation: Insights from Computational Modeling"

*Running title*: Modeling sex differences in AF mechanisms and therapy

Correspondence:

Eleonora Grandi, Department of Pharmacology, University of California Davis, 451 Health Science Drive, GBSF 3503, Davis, CA 95616, USA


Haibo Ni, Department of Pharmacology, University of California Davis, 451 Health Science Drive, GBSF 3503, Davis, CA 95616, USA


### Supplementary Tables

**Supplementary Table S1. Parameters for building sex-specific models of human atrial cardiomyocytes in nSR and cAF conditions.** $P_{ca}$ is the maximum of LTCC unitary current density, $k_{1o}$ regulates the LTCC $Ca^{2+}$-dependent inactivation rate, $k_{up}$ is $[Ca^{2+}]_{Cyto}$ for SERCA half-maximum flux, $K_{RyR,SR}$ is $[Ca^{2+}]_{SR}$ at half-maximum rate of RyR2 opening, and $K_b$ describes RyR2 sensitivity to $[Ca^{2+}]_{Cyto}$. $V_{up}$ depicts the maximum flux rate of SERCA.

| Parameters | nSR | | cAF | |
|---|---|---|---|---|
| | **Male** | **Female** | **Male** | **Female** |
| **Cell length (μm)** | 101.2 | 92.0 | 112.2 | 101.2 |
| **Cell width (μm)** | 15.3 | 11.7 | 18.0 | 14.4 |
| **Tubular density (μm/μm²)** | 0.542 | 0.274 | 0.013 | 0.008 |
| **LTCC unitary $Ca^{2+}$ permeability, $P_{Ca}$ ($\mu M \cdot C^{-1} \cdot ms^{-1}$)** | 8.188 | 8.188 | 4.094 | 8.188 |
| **LTCC $Ca^{2+}$-dependent inactivation rate, $k_{1o}$ ($ms^{-1}$)** | 0.104 | 0.104 | 0.034 | 0.104 |
| **CSQ expression, $B_{max}$ (μM)** | 400 | 304 | 220 | 220 |
| **SERCA sensitivity to $[Ca^{2+}]_{Cyto}$, $K_{up}$ (μM)** | 0.615 | 0.615 | 0.554 | 0.615 |
| **SR content threshold for RyR2 release, $K_{RyR,SR}$ (μM)** | 900 | 900 | 900 | 800 |
| **RyR2 sensitivity to $[Ca^{2+}]_{Cyto}$, $K_b$ ($ms^{-1}$)** | $5e^{-3}$ | $5e^{-3}$ | $5e^{-3}$ | $1.5e^{-2}$ |
| **Max. SERCA uptake rate, $V_{up}$ ($\mu M \cdot ms^{-1}$)** | 3.6 | 3.6 | 3.6 | 3.6 |

**Supplementary Table S2. Parameters describing the changes for each putative $Ca^{2+}$-directed intervention.**

| $Ca^{2+}$-directed interventions in cAF | Modified Parameter | Male | Female |
|---|---|---|---|
| **Tubular Up (restoration)** | ↑ Tubular density (μm/μm²) | 0.67 | 0.67 |
| **RyR2 Down (dephosphorylation)** | ↑ SR content threshold for RyR release, $K_{RyR,SR}$ (μM) | 1100 | 1100 |
| | ↓ RyR sensitivity to $[Ca^{2+}]_{Cyto}$, $K_b$ ($ms^{-1}$) | $2.5e^{-3}$ | $2.5e^{-3}$ |
| **SERCA Down (inhibition)** | ↓ SERCA strength of uptake, $V_{up}$ ($\mu M \cdot ms^{-1}$) | 1.8 | 1.8 |
| **CSQ Up (overexpression)** | ↑ CSQ expression, $B_{max}$ (μM) | 480 | 480 |
| **PLB Up (overexpression)** | ↑ SERCA $K_m$ value to $[Ca^{2+}]_{Cyto}$, $K_{up}$ (μM) | 0.738 | 0.738 |

#### Supplementary Figures

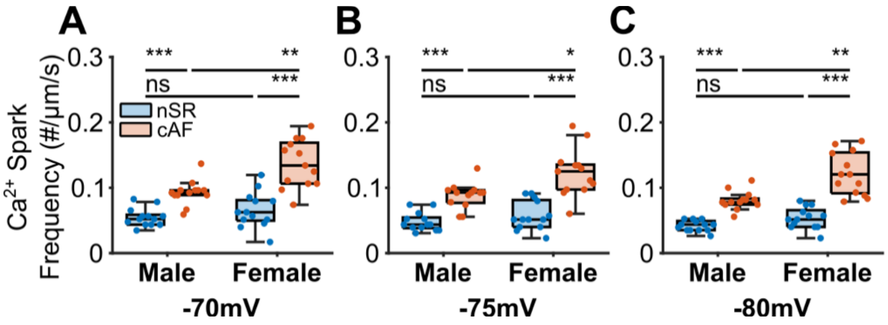


**Supplementary Fig. S1. Simulated sex differences of $Ca^{2+}$ sparks with AF conditions are independent of the holding membrane potential.** (**A-C**): the holding potential $V_{hold}$ = -70, -75, and -80 mV, respectively. For each group, data are reported from the analysis of 13 line scans in 1 cardiomyocyte model simulation. Statistical significance was determined by One-way ANOVA with planned comparison and Bonferroni correction and indicated with ***: $p < 0.001$; **: $p < 0.01$; *: $p < 0.05$; ns: not significant.

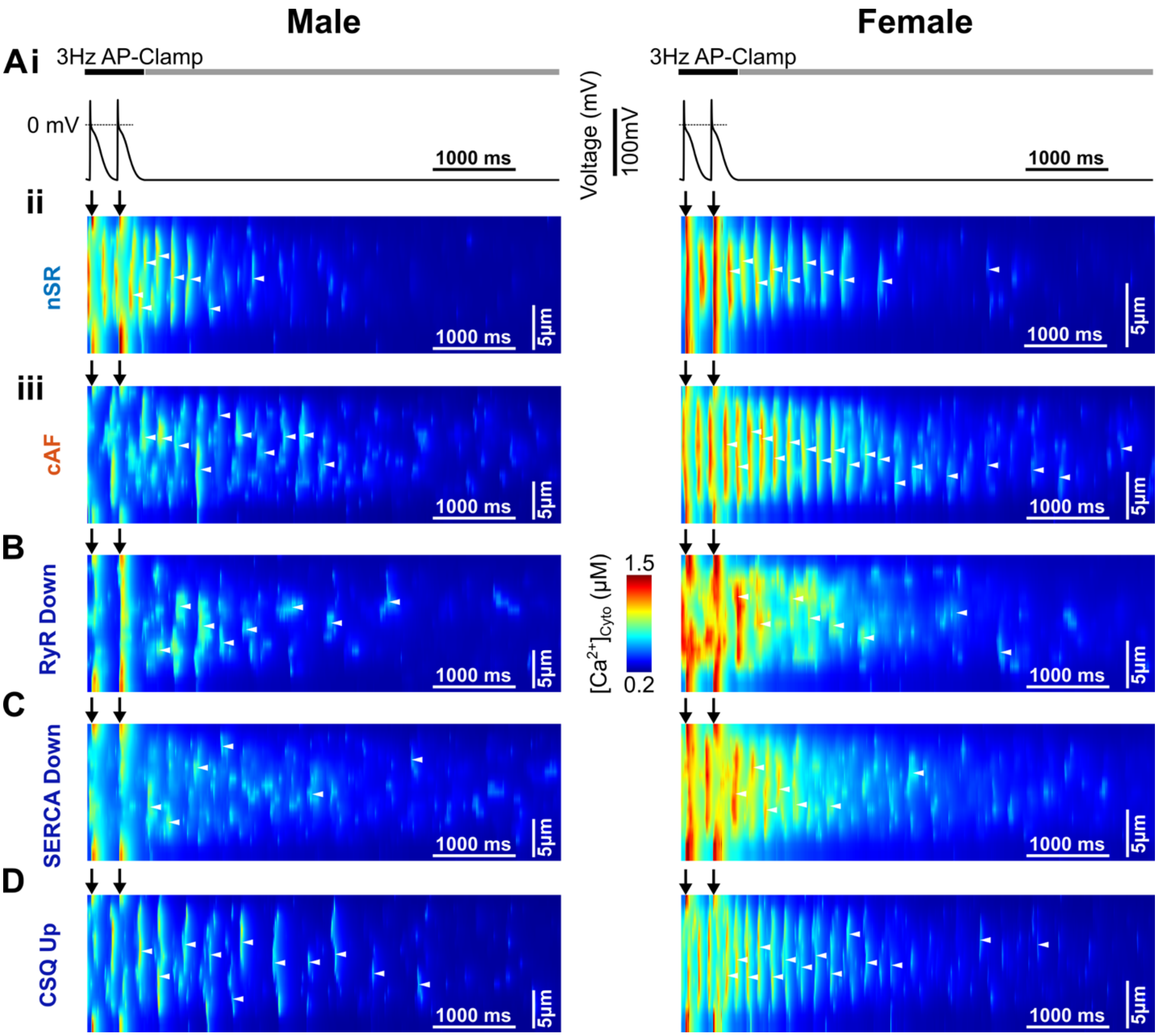


**Supplementary Fig. S2. Simulated effects of downregulating RyR2 and SERCA activities, and upregulating CSQ in AF-remodeled atrial cardiomyocytes**. **(A) (i)** AP-clamp traces, **(ii-iii)** simulated transversal line scans of $[Ca^{2+}]_{Cyto}$ for male and female cardiomyocytes in **(ii)** nSR and **(iii)** AF conditions as in **Fig. 2A** are illustrated for comparison. **(B-D)** Simulated transversal line scans of $[Ca^{2+}]_{Cyto}$ of human atrial cardiomyocytes in AF with interventions of downregulating **(B)** RyR2 and **(C)** SERCA activities, and **(D)** upregulating CSQ expression. White arrow heads indicate SCRs and black arrows mark the start of APs. For each group, data are reported from 1 cardiomyocyte model simulation.

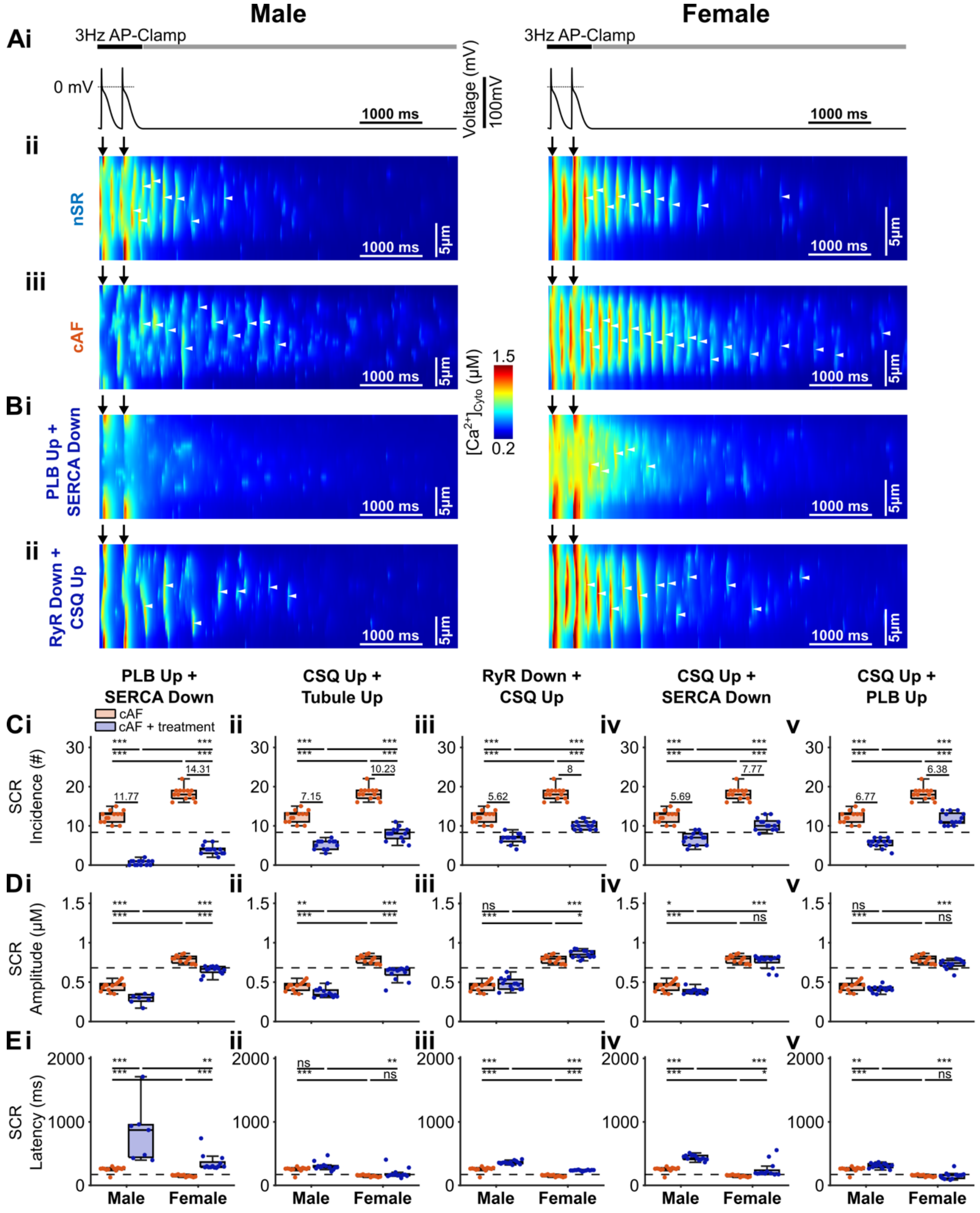


**Supplementary Fig. S3. Simulated effects of pair-wise combined interventions on male and female atrial cardiomyocytes from AF patients**. **(A)** **(i)** AP-clamp traces, **(ii-iii)** simulated transversal line scans of $[Ca^{2+}]_{Cyto}$ for male and female cardiomyocytes in **(ii)** nSR and **(iii)** AF conditions as in **Fig. 2A** are illustrated for comparison. **(B)** Simulated transversal line scans of $[Ca^{2+}]_{Cyto}$ with combined interventions of **(i)** PLB up + SERCA down and **(ii)** RyR2 down + CSQ up. White arrow heads indicate SCRs and black arrows mark the start of AP. **(C-E)** Summary of intervention effects on **(C)** SCR incidence, **(D)** SCR amplitude, and **(E)** SCR latency; columns **(i-v)** depict effects of combined interventions with (i) PLB up + SERCA down, (ii) CSQ up + Tubule up, (iii) RyR2 down + CSQ up, (iv) CSQ up + SERCA down, and (v) CSQ up + PLB up. In Panel (**C**), numbers indicate the mean reduction of SCR incidence due to intervention. For each group in (**C-E**), data are reported from the analysis of 13 line scans in 1 cardiomyocyte model simulation. Statistical significance was determined by One-way ANOVA with planned comparison and Bonferroni correction and indicated with ***: $p < 0.001$; **: $p < 0.01$; *: $p < 0.05$; ns: not significant.

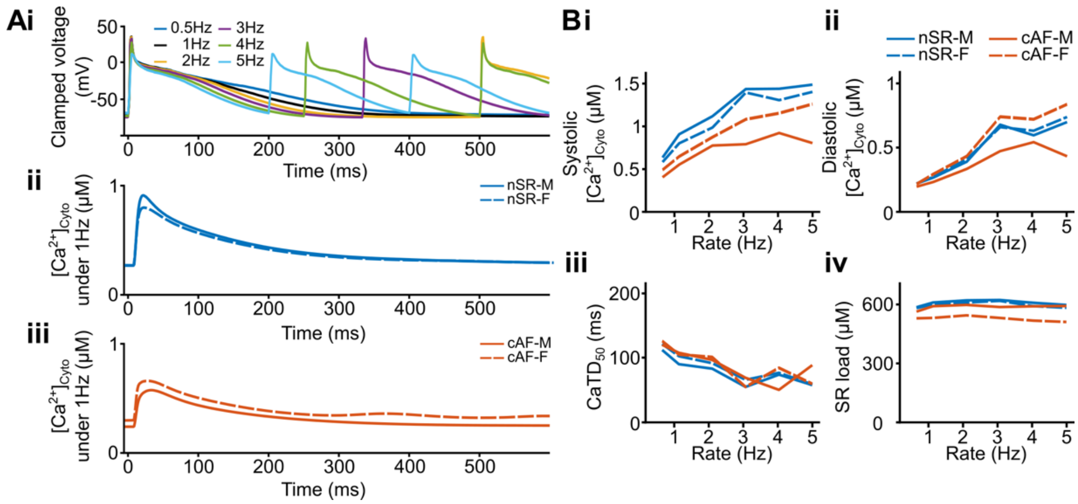


**Supplementary Fig. S4. Comparison of male (M) and female (F) intracellular $Ca^{2+}$ transient properties obtained with AP clamp repeated at 0.5 to 5 Hz. (A) (i)** Time courses of AP clamp traces at for 0.5 Hz to 5 Hz, and whole-cell averaged CaT in (**ii**) nSR and (**iii**) cAF conditions. **(B)** Rate-dependence of whole-cell averaged **(i)** systolic $[Ca^{2+}]_{Cyto}$, **(ii)** diastolic $[Ca^{2+}]_{Cyto}$, **(iii)** CaT duration at 50% recovery ($CaD_{50}$), and **(iv)** SR load. For each group, data are reported from 1 cardiomyocyte model simulation.

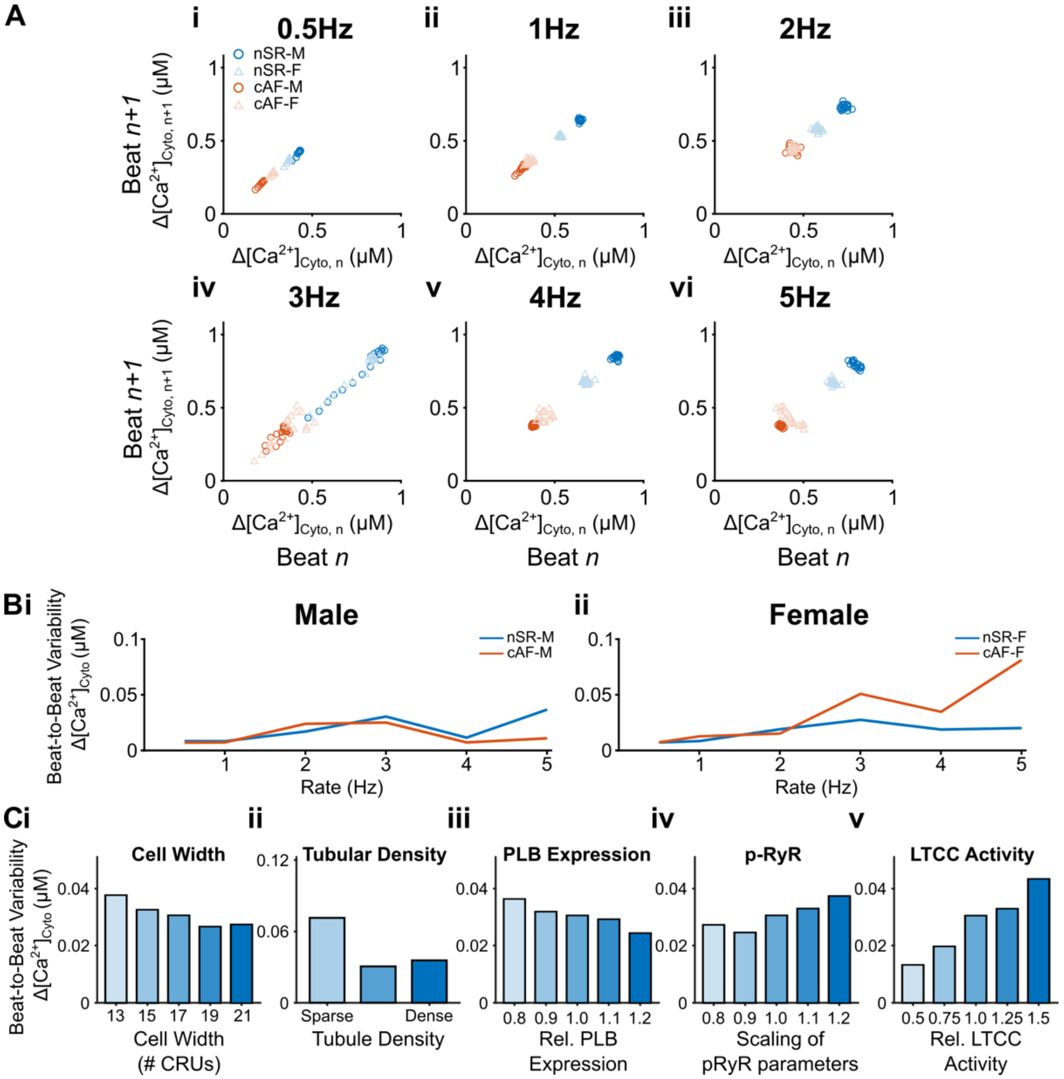


**Supplementary Fig. S5. Beat-to-beat variability of AP-clamp elicited CaTs. (A)** Poincaré plot illustrating the relation of the amplitudes of AP-clamp elicited CaT between beat *n* and beat (*n+1*) for AP-clamp repeated at multiple rates from (**i**) 0.5 Hz to (**vi**) 5 Hz. **(B)** Computed beat-to-beat variability of the amplitudes of AP-clamp evoked CaT for (**i**) male and (**ii**) female cardiomyocytes. The beat-to-beat variability was calculated as the mean absolute difference in the $Ca^{2+}$ transient amplitude between two consecutive beats. **(C)** Sensitivity analysis of beat-to-beat variability of CaT amplitude by varying (i) cell width, (ii) tubular density, (iii) PLB expression, (iv) RyR phosphorylation, and (v) LTCC activity. For each group, data are reported from 1 cardiomyocyte model simulation.

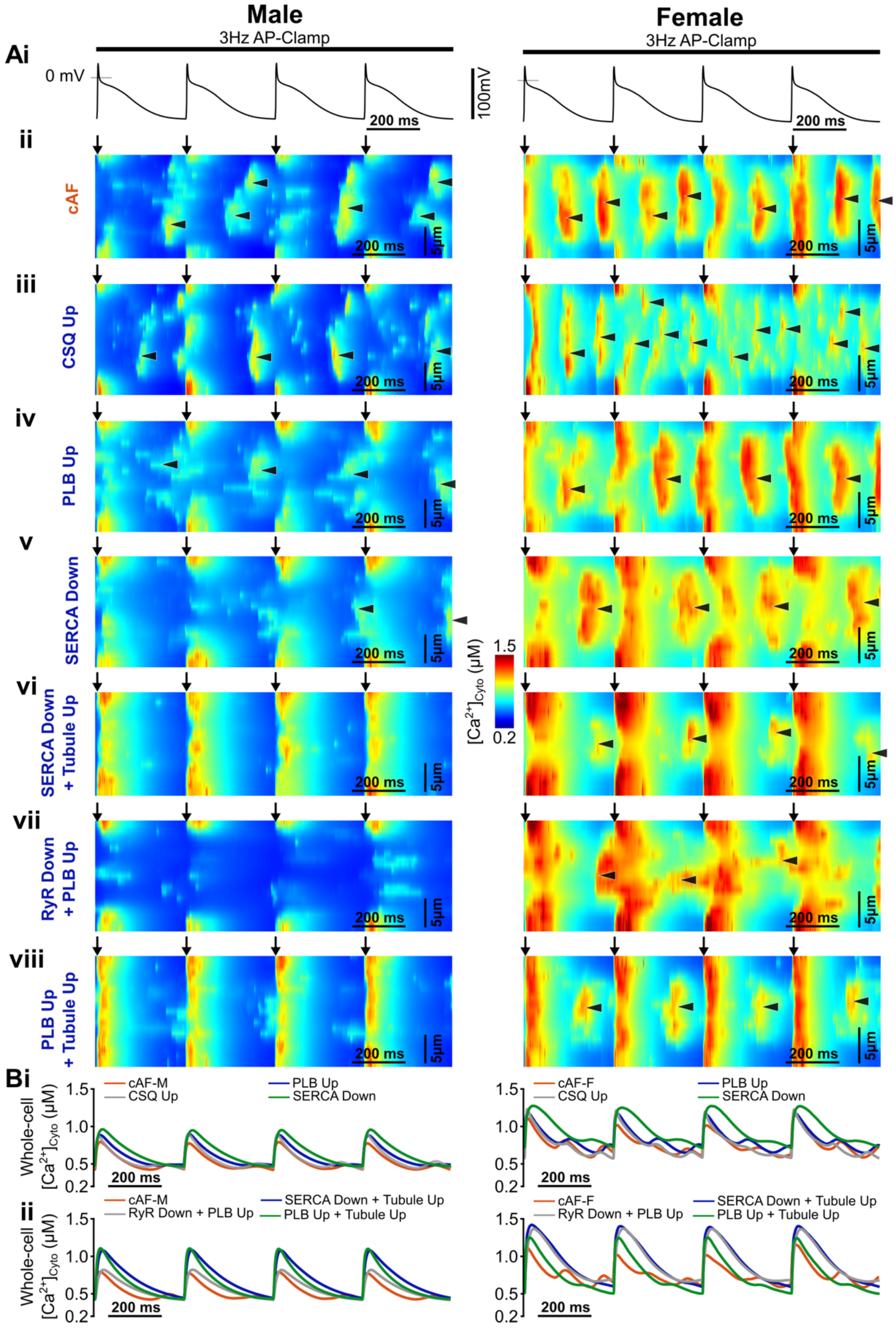


**Supplementary Fig. S6. Simulated transversal line scans of AP-clamp evoked $Ca^{2+}$ transients in male and female atrial cardiomyocytes and responses to interventions in AF.**

**(A)** **(i)** AP-clamp traces, **(ii-viii)** simulated transversal line scans of AP clamp-evoked $Ca^{2+}$ transients for male and female cardiomyocytes in **(ii)** AF conditions without intervention, and following interventions of **(iii)** CSQ up, **(iv)** PLB up, **(v)** SERCA down, and combined interventions of **(vi)** SERCA down + Tubule up, **(vii)** RyR2 down + PLB up, and **(viii)** PLB up + Tubule up. **(B)** Comparison of whole-cell averaged CaTs from cardiomyocytes in AF before intervention and with **(i)** single-hit and **(ii)** double-hit interventions. Arrows mark the start of APs, and the diastolic SCRs are indicated with arrow heads. For each group, data are reported from 1 cardiomyocyte model simulation.

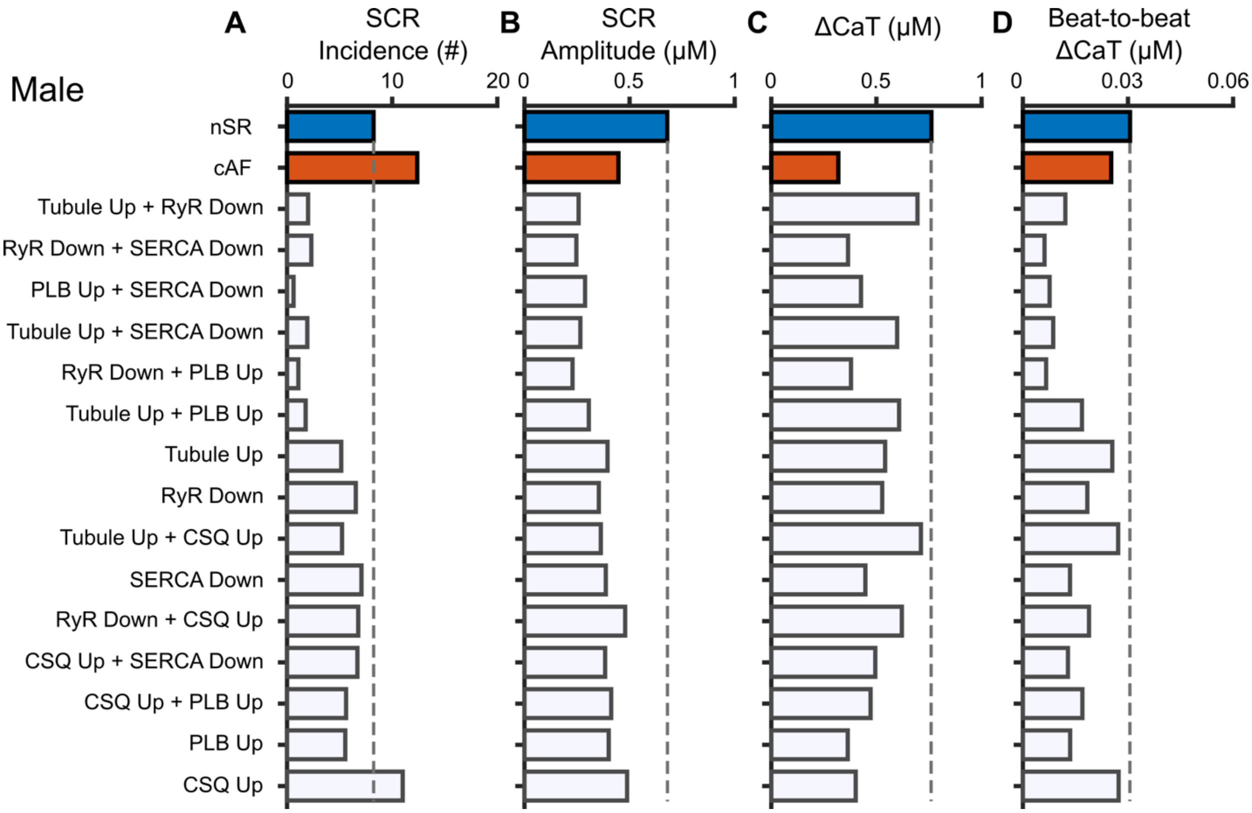


**Supplementary Fig. S7. Summary data comparing the intervention effects on SCR (A) incidence and (B) amplitude, (C) whole-cell CaT amplitude, and (D) CaT beat-to-beat variability in male atrial cardiomyocytes.** Dashed lines indicate the referencing values obtained from male cardiomyocytes in nSR conditions.
